# The RNA binding protein, HNRNPA2B1, regulates IFNG signaling in macrophages

**DOI:** 10.1101/2023.10.12.562050

**Authors:** Mays Mohammed Salih, Chi G Weindel, Eric Malekos, Lisa Sudek, Sol Katzman, Cory J Mabry, Aja K Coleman, Sikandar Azam, Robert Watson, Kristin Patrick, Susan Carpenter

## Abstract

Heterogeneous nuclear ribonucleoprotein A2B1 (HNRNPA2B1) is a well known RNA binding protein but the mechanisms by which it contributes to innate immune gene regulation are poorly understood. Here we report that HNRNPA2B1 functions in macrophages to regulate IFNG (IFN-γ) signaling through alternative splicing of the IFNG receptor. Specific deletion of HNRNPA2B1 in macrophages resulted in altered cytokine responses in both an endotoxic shock model and following *Salmonella* infection. Interestingly, while HNRNPA2B1 can function as a viability gene, we observed increased macrophage and neutrophil numbers in the KO mice following LPS induced endotoxic shock. We also discovered that HNRNPA2B1 restricts replication of *Salmonella enterica in vivo*. Mechanistically, loss of HNRNPA2B1 resulted in an increase in NGO transcripts, which lack a start codon, of the IFNG receptor (*Ifngr*) leading to lower expression of the receptor at the cell surface impacting the downstream IFNG signaling cascade. Collectively, our data highlight an important role for HNRNPA2B1 in regulating IFNG signaling and restricting intracellular bacterial pathogens in macrophages.

## Introduction

Inflammation is the host’s natural response to protect against infection and maintain homeostasis. Macrophages are crucial components of the innate immune system required for mediating the magnitude and outcome of the inflammatory response. By illuminating the mechanisms through which macrophages regulate this response, we will be better equipped to combat infection and inflammatory diseases. Macrophages sense danger signals through PRRs (1,2) -such as TLR4-which recognizes lipopolysaccharide (LPS) triggering complex signaling cascades culminating in the production of pro-inflammatory cytokines (1,3–5). Due to the complex nature of these responses there are regulatory steps throughout including at the level of transcription, pre-mRNA splicing, RNA modification, RNA export, and translation (6). In recent years a small number of studies have been carried out to investigate the role that HNRNP proteins play in regulating innate immune responses. It is well documented that HNRNPs represent ubiquitously expressed proteins critical for RNA processing and yet some members of the family play highly specific roles in certain cell types (7). HNRNPM regulates gene expression through repression of splicing in macrophages. Repression is only relieved on target genes such as *Il6* following phosphorylation of HNRNPM (8). HNRNPU has been reported to translocate from the nucleus to the cytosol following LPS stimulation in macrophages leading to the stabilization of target mRNA including *Tnf* and *Il6* (9). HNRNPA0 has also been implicated in regulating mRNA within the cytosol of macrophages following LPS stimulation, by interacting with AU rich elements in target genes such as TNF influencing their expression (10).

HNRNPA2B1 is a highly conserved, abundant member of the HNRNPA/B subfamily which is involved in all aspects of RNA metabolism from biogenesis to degradation (e.g. processing and splicing, trafficking, mRNA translation and stability) (11, 12). HNRNPA2B1 is localized to the nucleus (12, 13) allowing it to mediate gene expression regulation through its control over transcription initiation (14, 15, 16), alternative splicing (17–24) and RNA export (25–27). HNRNPA2B1 plays a central role in RNA metabolism and dysregulation and mutations in this protein are associated with a range of metabolic and neurodegenerative diseases as well as cancers (17, 28–31). Recently HNRNPA2B1 has been reported to function as a DNA sensor within the nucleus facilitating interferon signaling downstream of HSV-1 infection (32). Importantly, loss of tolerance to HNRNPA2B1 is recognized as a hallmark of several systemic autoimmune rheumatic diseases (SARDs) such as systemic lupus erythematosus (SLE) and rheumatoid arthritis (RA) (33–37).

While our group and others have previously shown that HNRNPA2B1 can play a role in regulating the innate immune response downstream of TLR signaling *ex vivo* (15, 38), we do not understand the mechanisms by which HNRNPA2B1 regulates gene expression *in vivo*. To answer this question, we generated a conditional mouse and depleted HNRNPA2B1 in myeloid cells by crossing to *LysM*Cre. RNA-seq was carried out following LPS stimulation comparing CTL (HNRNPA2B1^fl/fl^) and KO (HNRNPA2B1^fl/fl^ *LysM*Cre^+/+^) macrophages. While many genes were altered in expression at baseline and following stimulation in the knockout, one of the most impacted signaling cascades was that of interferon gamma (IFNG) with reduced expression of key components from receptors (IFNGR) to signaling components (STATs), as well as transcription factors (IRFs). Interestingly the dampened IFNG response resulted in mice being less responsive to LPS induced endotoxic shock but more susceptible to *Salmonella* infection, wherein loss of HNRNPA2B1 led to uncontrolled bacterial replication. Mechanistically, loss of HNRNPA2B1 resulted in an increase in NGO transcripts of the IFNG receptor1 (*IfngrI*) leading to lower expression of the receptor at the cell surface impacting the entire downstream IFNG signaling cascade. Collectively our work provides insights into HNRNPA2B1 as a specific regulator of the IFNG signaling cascade in macrophages.

## Results

### The immune cell repertoires of HNRNPA2B1 knockout mice are phenotypically normal at steady state

Due to HNRNPA2B1’s wide ranging roles in RNA metabolism it is considered a viability gene, consistent with our failed attempts to generate a full body knockout of HNRNPA2B1. Therefore, to investigate HNRNPA2B1 function in innate immune cells, we generated a HNRNPA2B1 conditional knockout (KO) mouse using a *LysM*Cre system (Fig.1, A) (SFig. 1). We inserted two loxP sites flanking exons 2 and 7 of the HNRNPA2B1 locus using CRISPR and crossed to *LysM*Cre to homozygosity to selectively delete HNRNPA2B1 from myeloid cells. Western blot analysis confirmed complete knockout of HNRNPA2B1 in the deficient bone marrow derived macrophages (BMDMs) (Fig.1, B). Mice appeared normal and bred at expected mendelian ratios. Immune populations were profiled at baseline in blood and spleen and no difference was observed in immune cell numbers for monocytes, macrophages, eosinophils, T or B cells (Fig.1, C and D). A small increase in neutrophils was recorded in the spleen (Fig. 1D). HNRNPA2B1 deletion impaired macrophage *ex vivo* proliferation at day 7 post differentiation consistent with our previous CRISPR screen findings showing it acting as a viability gene in macrophages *ex vivo* (39), however, it did not impact macrophage phagocytic function as measured by pHrodo Green *E. coli* particle uptake (Fig.1, E and F).

**Figure 1.**
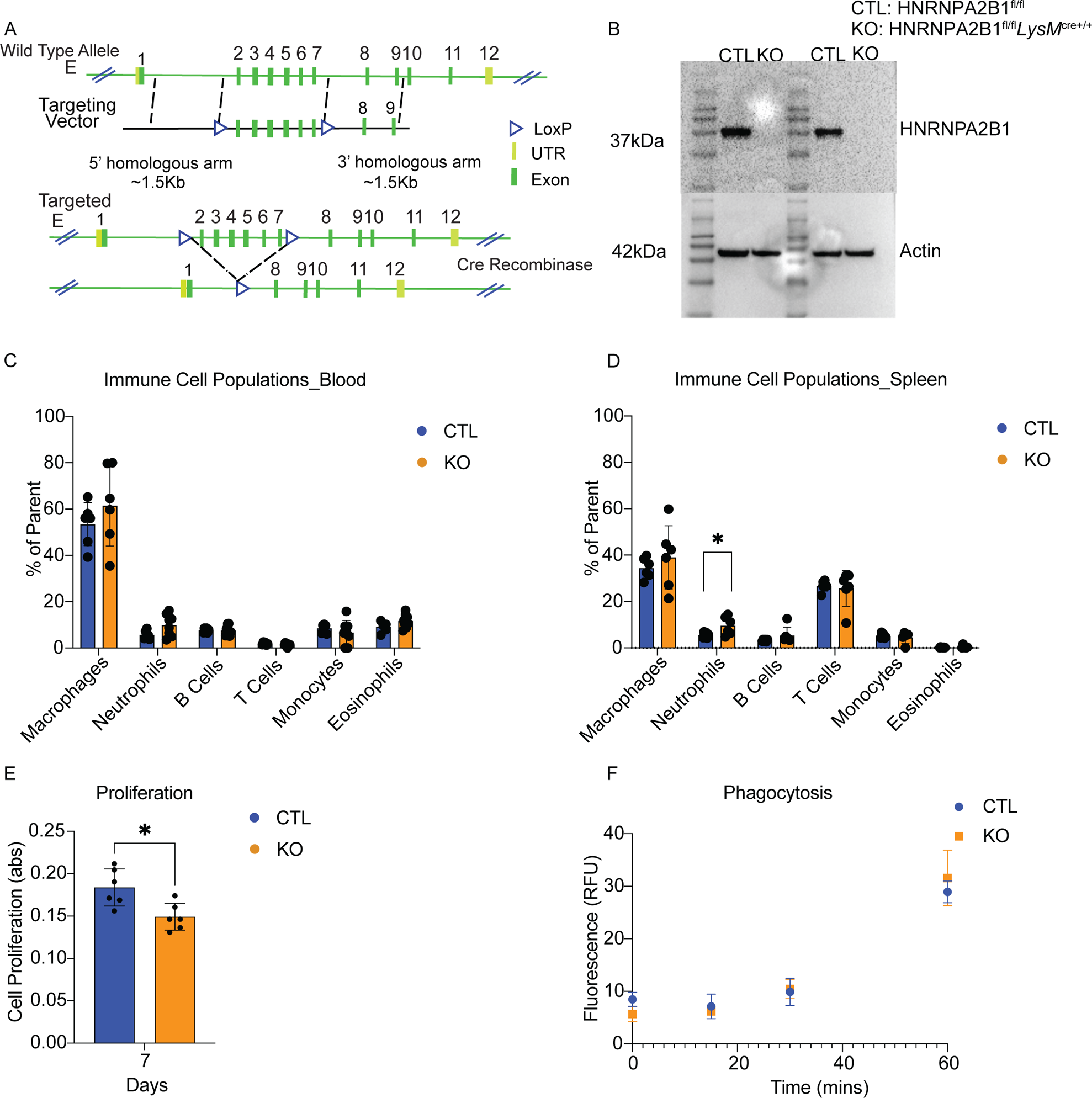
Generation and characterization of the HNRNPA2B1 conditional knockout mouse. **A.** CRISPR was used to add loxP sites on the 5’ and 3’ ends around exons 2 and 7 of the HNRNPA2B1 locus. **B.** Western blot analysis of HNRNPA2B1 levels in murine BMDMs. **C.** Profiling of the immune cell repertoire in the blood of HNRNPA2B1 KO mice. **D.** Profiling of the immune cell repertoire in the spleen of HNRNPA2B1 KO mice. **E.** MTT proliferation assay was performed to assess BMDM proliferation rate in HNRNPA2B1 KO BMDMs. **F.** PHrodo green *E. coli* bioparticle assay was used to assess BMDM phagocytic function.

### IFN signaling cascades are downregulated in HNRNPA2B1 KO macrophages in response to LPS

In order to determine what role HNRNPA2B1 plays in regulating gene expression in macrophages we first performed RNA sequencing on KO BMDMs compared to CTL at baseline and following 5 h of LPS stimulation. Differential expression analysis (DESeq2) revealed a large number of upregulated (178) and downregulated (66) genes in macrophages at baseline, as well as 204 upregulated and 98 downregulated genes under inflammatory conditions when HNRNPA2B1 is knocked out (Fig. 2, A and B). Gene ontology (GO-term) analysis of differentially expressed genes revealed that HNRNPA2B1 loss led to an overall downregulation of inflammatory response genes such as *Cd74* and *Mill2* at baseline, as well as genes such as *Irf8,* and *Cxcl9* after LPS treatment (Fig. 2, A, B, C and E). Upregulated GO-terms at baseline consisted mostly of genes involved in cell cycle and cell division regulation as well as DNA replication (Fig. 2, D and F). Among a large number of downregulated genes in LPS-treated BMDMs, there was a strong enrichment of interferon response genes, specifically those involved in the IFNG (IFN-γ) response signaling pathway such as *IfngrII, Gbp2* and *Cd74* (Fig. 2, G). Using RT-qPCR, we confirmed lower expression levels in KO BMDMs of several major IFN response genes (*Irf7, Irf8, Stat3,* and *Oas1c*) and an increase in *Ifi208* (SFig. 2, A-E). To determine the impact these transcriptional changes, have on the downstream IFNG cascade we utilized western blot analysis to assess changes in phosphorylation of STAT3, which is a major transcription factor (TF) in the IFNG signaling pathway. Our results revealed that phosphorylated STAT3 (pSTAT3) was activated 2 h and 5 h post LPS stimulation in CTL mice, but this was significantly reduced in the KO BMDMs (Fig. 2, H). Total STAT3 was similar between CTL and KOs (Fig. 2, H). Using multiplex ELISA, we measured altered expression for many key inflammatory cytokines in HNRNPA2B1 KO BMDMs, with some showing increased expression (CXCL5, CCL22, CSF1, CSF3, CCL17, IL1B, IL5, IL12, and IL15), others showing decreased expression (e.g. CXCL9, IL10, and TIMP1) post-stimulation (Fig.2, I-L, SFig. 3, A-H), and a number of proteins remained unchanged (SFig. 3, I-M). Collectively, our data indicate that in macrophages LPS induced cytokine responses are altered when HNRNPA2B1 is removed and many components of the IFN signaling pathway are downregulated.

**Figure 2.**
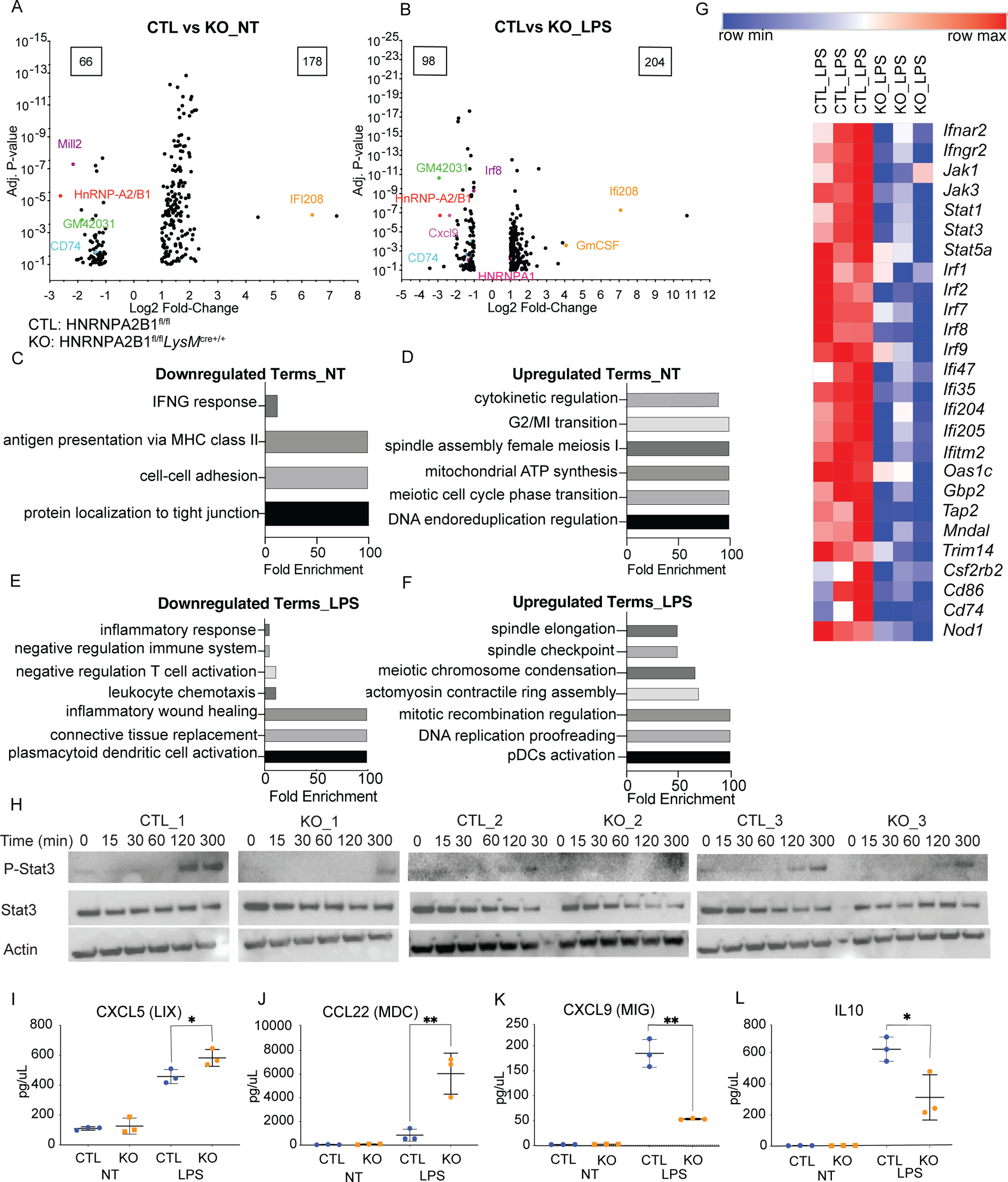
HNRNPA2B1 KO macrophages have altered IFN signaling in response to LPS. **A.** Volcano plot of differentially expressed genes in HNRNPA2B1 KO BMDMs at baseline. **B.** Volcano plot of differentially expressed genes in HNRNPA2B1 KO BMDMs under LPS stimulation. **C.** Gene ontology analysis of downregulated genes at baseline. **D.** Gene ontology analysis of upregulated genes at baseline. **E.** Gene ontology analysis of downregulated genes post LPS stimulation. **F.** Gene ontology analysis of upregulated genes post LPS stimulation. **G**. Heat map of IFNG response genes normalized counts in CTL and KO cells post LPS stimulation. **H.** Western blot analysis of pSTAT3 and STAT3 levels in HNRNPA2B1 KO BMDMs under stimulus (right panel). **I-L.** Cytokine levels as measured by ELISA from BMDM supernatant, the supernatant was harvested from CTL and KO cultured BMDMs and multiplex cytokine analysis was performed for (I) CXCL5 (LIX), (J) CCL22 (MDC), (K) CXCL9 (MIG), (L) IL10. Each dot represents BMDMs from an individual animal. Error bars represent the standard deviation of biological triplicates. Student’s t-tests were performed using GraphPad Prism. Asterisks indicate statistically significant differences between mouse lines (*P ≤ 0.05, **P ≤ 0.01).

### HNRNPA2B1 KO mice display altered immune responses following endotoxic shock *in vivo*

Next, we investigated whether the observed disruption in macrophage responses would lead to altered responses to endotoxic shock *in vivo* in the KO mice. One early clinical feature of endotoxic shock in mice is the rapid decrease in body temperature. Here we recorded an average temperature of 25°C for the CTL mice while the HNRNPA2B1 KO mice were at ∼32°C following 5 mg/kg intraperitoneal injection of LPS for 18 h (Fig.3, A). In addition, cytokine analysis of serum, spleen and liver revealed an impaired cytokine production represented by the lower inflammatory cytokine levels in the KO mice (Fig.3, B-N). Most importantly we observed reduced levels of IFNG in the serum of KO mice (Fig. 3, B) consistent with what we observed in the macrophage *ex vivo* experiment in Fig. 2. Some impaired cytokines were common in serum and spleen (e.g. CCL2 and CCL3) (Fig.3, C, D, H and I), while others were reduced in serum only (IFNG, CSF1 and CCL5) (Fig. 3, B, E and F) or were restricted to the spleen (IL6, CSF2 and CXCL1) (Fig.3, G, J and K). Only IL20 was reduced in the KO livers compared to CTL while VEGF and TIMP1 were increased (Fig. 3, M, and N). From these data we can conclude that specific loss of HNRNPA2B1 from myeloid cells resulted in significant changes in the pro-inflammatory cytokine levels throughout the tissues of the knockout mice indicating that HNRNPA2B1 plays an important and specific role in regulating gene expression in these cells.

**Figure 3.**
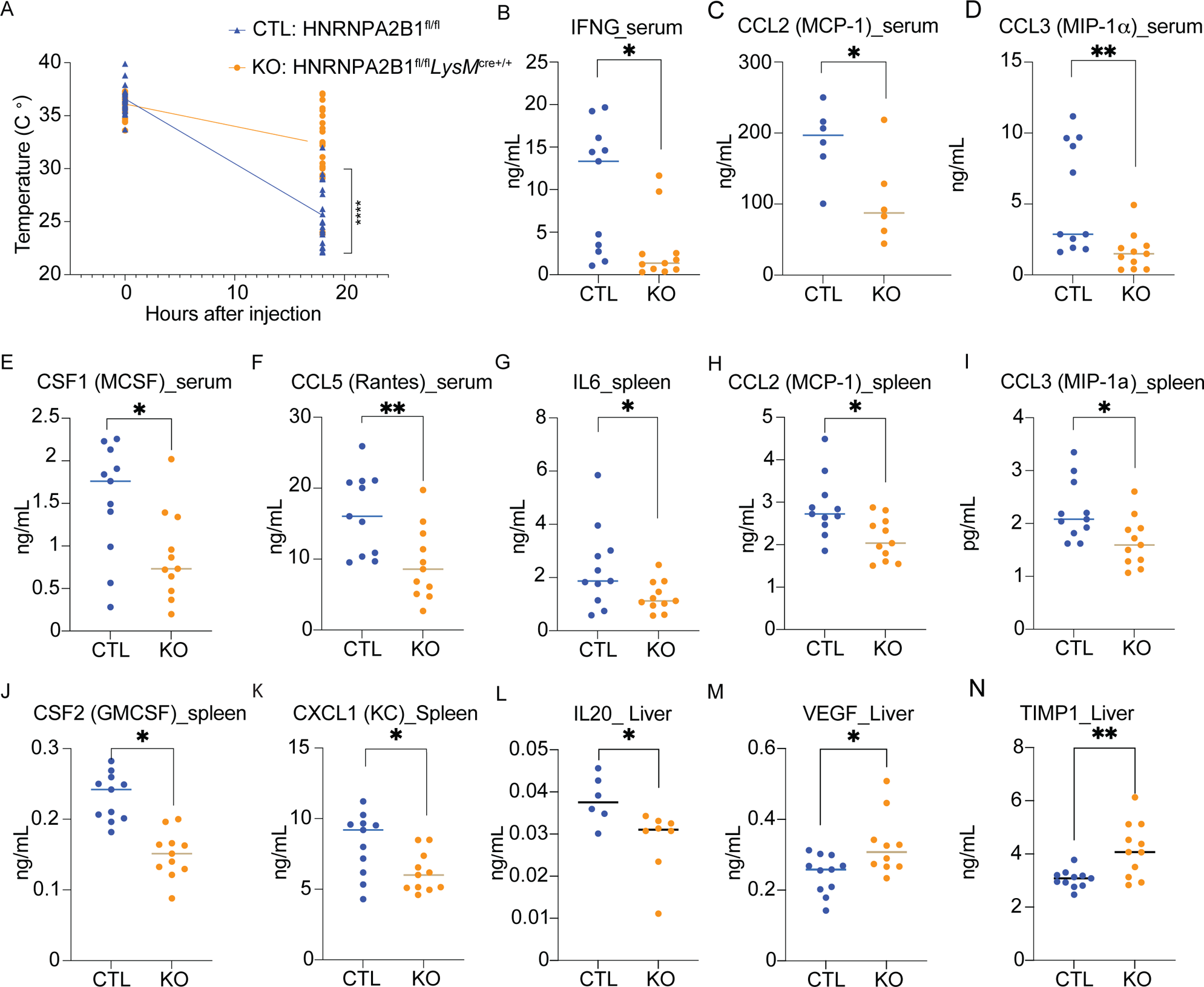
HNRNPA2B1 KO mice show altered immune responses following endotoxic shock *in vivo*. **A.** Temperature change in CTL and HNRNPA2B1 KO mice after i.p. Injection with 5 mg/kg LPS. N= 25 CTL, 26 KO. **B-N.** Cytokine levels in serum, spleen and liver of mice treated with 5 mg/kg LPS 6 hrs post injection. **(B)** IFNG_serum, **(C)** CCL2_serum, **(D)** CCL3_serum, **(E)** CSF1_serum, **(F)** CCL5_serum, **(G)** IL6_spleen, **(H)** CCL2_spleen, **(I)** CCL3_spleen, **(J)** CSF2_spleen, **(K)** CXCL1_spleen, **(L)** IL20_Liver, **(M)** VEGF_Liver, **(N)** TIMP1_Liver. Student’s t-tests were performed using GraphPad Prism. Asterisks indicate statistically significant differences between mouse lines (*P ≤ 0.05, **P ≤ 0.01).

### Macrophage and neutrophil numbers are elevated in the HNRNPA2B1 KO mice and macrophages display altered costimulatory molecule expression

Since HNRNPA2B1 is a known viability gene we speculated that the downregulated responses could be simply due to less macrophages and neutrophils in the KO mice. Interestingly we found the opposite to be true with the KOs showing increased macrophage and neutrophil counts in the spleen and blood following endotoxic shock (Fig.4, A-D). There was no difference in other immune cells such as B cells, monocytes and eosinophils in the KO mouse, except for T cells that displayed a small increase in KO mouse spleen (SFig.4, A-H). We also investigated whether the observed elevation in macrophage numbers was extended to other macrophage subsets such as peritoneal macrophages (PMs) since they are directly exposed to LPS upon injection, but we observed no changes in PM levels post LPS introduction (SFig. 4, I and J).

**Figure 4.**
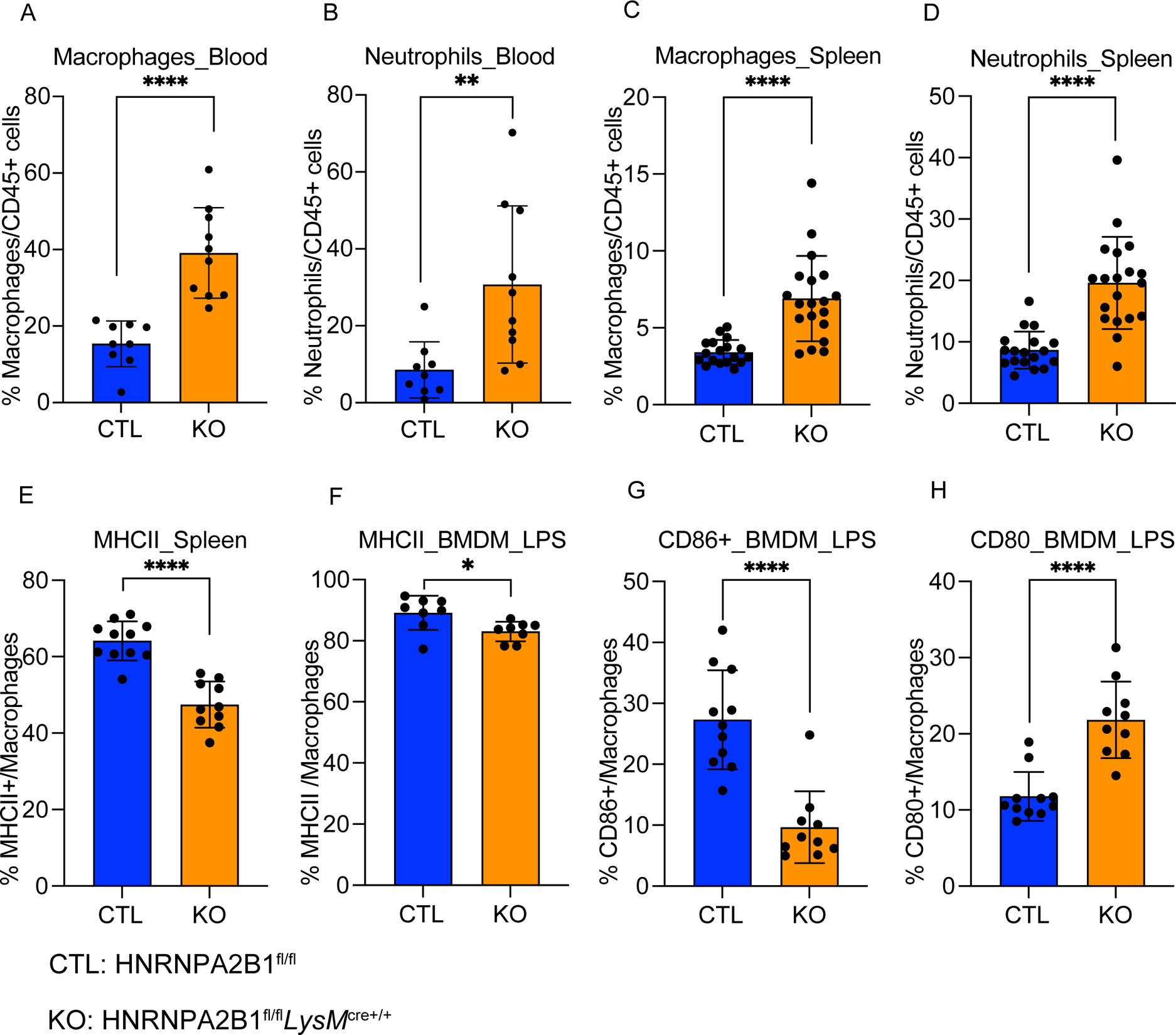
Macrophage and neutrophil levels are elevated in the HNRNPA2B1 KO mice and show altered macrophage activation following endotoxic shock. **A.** Macrophage levels from CTL and HNRNPA2B1 KO mice blood, measured using a flow cytometry panel. **B.** Neutrophil levels from CTL and HNRNPA2B1 KO mice blood, measured using a flow cytometry panel. **C.** Macrophage levels from CTL and HNRNPA2B1 KO mice spleen, measured using a flow cytometry panel. **D.** Neutrophil levels from CTL and HNRNPA2B1 KO mice spleen, measured using a flow cytometry panel. **E.** Level of MHCII marker on macrophage surface in the spleen, analysis performed using flow cytometry. **F-H** Level of activation markers on BMDMs *ex vivo*, analysis performed using flow cytometry **(F)** MHCII, **(G)** CD86, **(I)** CD80. Student’s t-tests were performed using GraphPad Prism. Asterisks indicate statistically significant differences between mouse lines (*P ≤ 0.05, **P ≤ 0.01, ****P ≤ 0.0001).

Macrophage activation is an important indicator of macrophage function and ability to respond to challenges. Thus, we assessed macrophage activation markers and found MHC Class II expression to be lower in the spleen, which we confirmed in macrophages *ex vivo* (Fig. 4, E and F). In addition, we found CD86 levels to be reduced while CD80 levels are higher in BMDMs following LPS stimulation (Fig. 4, G and H). Thus, HNRNPA2B1 KO macrophages are more abundant in LPS-exposed mice, however, they are less proinflammatory and display attenuated responses following inflammatory activation with LPS.

### HNRNPA2B1 regulates IFNG signaling through alternative splicing

HNRNPA2B1 is a well-recognized regulator of alternative splicing (21–24). Having confirmed strict nuclear localization of HNRNPA2B1 under a variety of inflammatory stimuli (Fig.5, A) (SFig.5, A), we hypothesized that it regulates interferon response genes through alternative splicing. We employed Isoform Usage Two-step Analysis (IUTA) (40) to detect differential usage of gene isoforms in the KO BMDMs after LPS treatment. We found that only a small portion of the downregulated (DE) genes were alternatively spliced (213 out of 1389) in the KO macrophage. Likewise, only 43 out of 1024 upregulated DE genes were alternatively spliced in HNRNPA2B1 KO macrophages (Fig.5, B). GO-term analysis revealed that downregulated alternatively spliced genes were involved in the inflammatory response, specifically, IFN response (Fig.5, C). Taking into consideration the changes in IFNG receptor (*Ifngr)* gene expression in the KO BMDMs (Fig. 2, G) and the lower level of IFNG cytokine production in the KO mouse (Fig.3, B), we speculated that these changes are modulated by alternative splicing events in the *Ifngr* locus. Our IUTA analysis revealed differential isoform expression in the *IfngrI* locus as a result of HNRNPA2B1 deletion, leading to increased expression of a NGO transcript, which we identified using FLAIR (41), that lacks a start codon and therefore is not translated (Fig.5, D). We quantified both IFNGRI and II levels in macrophages from mice post LPS injection (Fig. 5, E and G), and on BMDMs treated with LPS *ex vivo* (Fig. 5, F) and confirmed lower levels of the receptors on the cell surface. We also identified similar alternative splicing events leading to an increase in NGO transcripts of major IFN response transcription factors (TFs) and genes such as (*Stat3*, *Stat1*, *Irf7* and *Oas1c*) (SFig. 6, A-D). Our results indicate that HNRNPA2B1 modulates IFNG signaling through alternative splicing of the IFNGR transcript, which dampens IFNGR levels on the macrophage surface.

**Figure 5.**
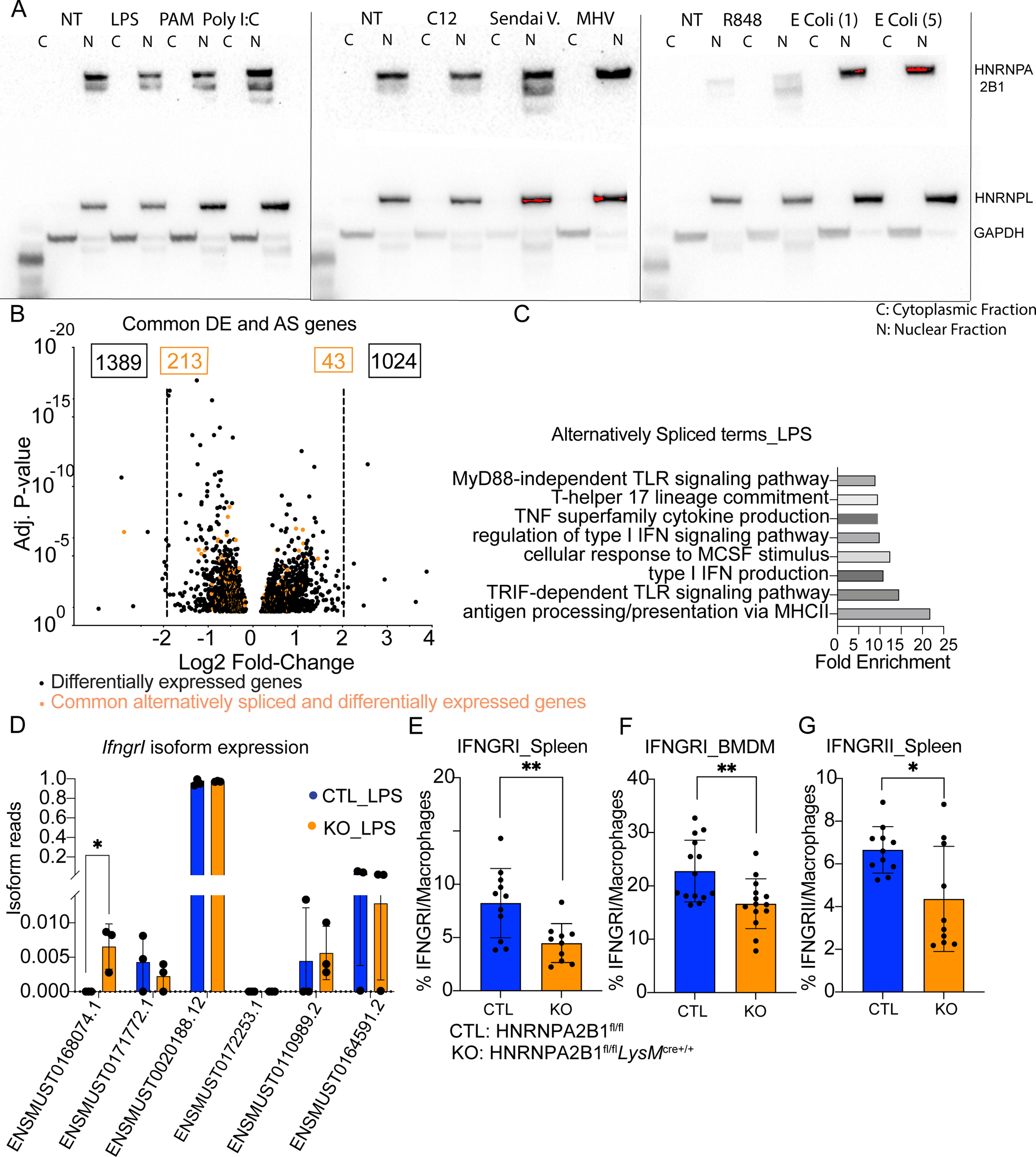
HNRNPA2B1 regulates IFNG signaling through alternative splicing. **A.** CTL BMDM nuclear and cytoplasmic fractions were analyzed using western blots after treatment with a panel of immune stimuli to assess changes in HNRNPA2B1 localization. **B.** Volcano plot of DE genes (black) and common DE and alternatively spliced genes in orange in KO BMDMs. **C.** Gene ontology analysis of genes that are differentially expressed as well as alternatively spliced in KO BMDMs. **D.** *IfngrI* isoform expression analysis performed using IUTA in CTL and KO BMDMs. **E.** IFNGRI levels on macrophages in the spleen using flow cytometry. **F.** IFNGRI levels on macrophages *ex vivo* using flow cytometry. **G.** IFNGRII levels on macrophages in the spleen using flow cytometry. Student’s t-tests were performed using GraphPad Prism. Asterisks indicate statistically significant differences between mouse lines (*P ≤ 0.05, **P ≤ 0.01).

### HNRNPA2B1 KO mice are susceptible to *Salmonella* infections due to a failure to clear the pathogen

Given the altered inflammatory responses observed in the HNRNPA2B1 KO mice during endotoxic shock, we wanted to investigate the role of HNRNPA2B1 in controlling the innate immune response to pathogens. To this end, we chose *Salmonella enterica* serovar Typhimurium as it utilizes macrophages as a replicative niche and is controlled in macrophages via IFNG (42). We asked whether loss of HNRNPA2B1 in macrophages impacted susceptibility to bacterial infection in an *in vivo* model. Briefly, we infected CTL and HNRNPA2B1 KO mice by IP delivery of *Salmonella* (2.5×10^4^) and mice were followed for signs of imminent morbidity (e.g. hunched posture, ruffled coat, lethargy as described in 43) over the course of 6 days. We observed that HNRNPA2B1 KO mice succumbed to infection earlier than controls (Fig. 6A) and experienced significantly higher bacterial burdens in the spleen and mesenteric lymph nodes (mLNs) (Fig. 6, B and C). Mouse susceptibility to infection was also represented by reduced expression of IL12, IL13, CXCL5, and CCL11 (Fig. 6, D-G). Our data indicate that HNRNPA2B1 KO mice are more susceptible to *Salmonella* infection and fail to produce sufficient levels of key pro-inflammatory cytokines in response to bacterial infection.

**Figure 6.**
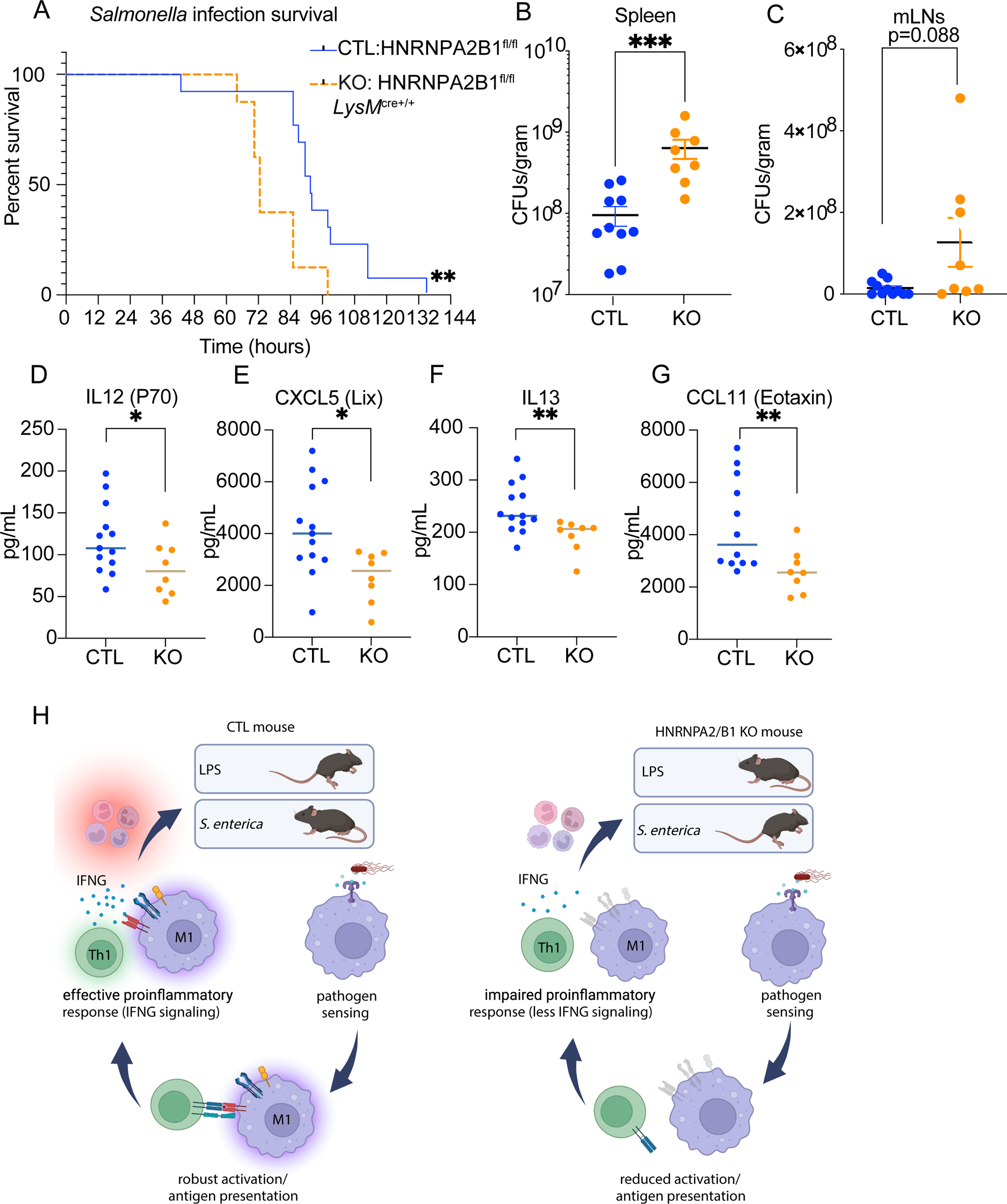
HNRNPA2B1 KO mice are susceptible to macrophage specific infection due to KO macrophage’s failure to clear the pathogen. **A.** Survival of CTL and HNRNPA2B1 KO mice after i.p. *Salmonella* infection. **B.** CFU measurement of *Salmonella* bacterial load in mouse spleen. **C.** CFU measurement of *Salmonella* bacterial load in mouse mesenteric lymph nodes. **D-G.** Cytokine levels in serum of CTL and KO mice infected with *Salmonella*. **(D)** Il12 (P70), **(E)** CXCL5, **(F)** IL13, **(G)** CCL11. Student’s t-tests were performed using GraphPad Prism. Asterisks indicate statistically significant differences between mouse lines (*P ≤ 0.05, **P ≤ 0.01).

## Discussion

HNRNPA2B1 is known to play critical roles in processes involving transcription, pre-mRNA splicing and translation (15,21–25,44–46) yet we find it is not essential for viability in macrophages or neutrophils. Using *LysM*Cre we generated a complete knockout in macrophages with only a slight decreased proliferative capacity observed *ex vivo*. HNRNPs have been shown to compensate for each other which is possibly occurring in the KO macrophages as our RNA-seq data indicated increased levels of HNRNPA1 in the HNRNPA2B1 deficient cells. It is possible that the importance of HNRNPA2B1 for macrophage fitness is also more important as cells age which could be the basis of future studies to determine. We were unsuccessful in our attempts to generate a HNRNPA2B1 knockout immortalized cell line using the CreJ2 method which supports the idea that over time HNRNPA2B1 is important for fitness as it takes upwards of 3 months of growth for cells to fully immortalize (data not shown, 47). We observed a widespread defect in cytokine production in the deficient mice following endotoxic shock. While we initially hypothesized that lower cytokine expression *in vivo* could be explained by reduced myeloid cell numbers given the lower *ex vivo* proliferative rate, surprisingly, macrophage numbers were higher in HNRNPA2B1 deficient mice. Elevated macrophage numbers were unexpected given that suppression of proliferation classically occurs upon exposure to proinflammatory stimuli such as LPS in conjunction with a switch to glycolysis in order to conserve the cell’s metabolic capacity (48,49). Despite being more abundant, KO macrophages were less active as evidenced by lower levels of M1 polarization markers (CD86 and MHCII) (50–53) in response to LPS. The increase in macrophage and neutrophil population numbers in the KOs could be a direct result of the weakened response to stimulus and failure to reach a terminally polarized state, prompting the cells to proliferate at a faster rate in order to attempt to compensate for the attenuated response.

We previously reported that HNRNPA2B1 regulates immune gene expression using shRNAs in macrophages *ex vivo* (15), where HNRNPA2B1 knockdown led to increased expression of a subset of IFN response genes. Here we performed RNA sequencing analysis comparing HNRNPA2B1 KOs to CTL and similar to shRNA mediated silencing, we observed widespread changes in the macrophage transcriptome. However, the effects we observed in the HNRNPA2B1 knockouts are more pronounced compared to the shRNA results likely due to more efficient removal of the gene *in vivo* and the use of primary macrophages rather than immortalized cells. While some genes were upregulated, the dominant phenotype was an overall dampening of the IFN response supported by down regulation in *Cd74* (54–56), *Mill2* (57), *MhcII* (H2Aa) (58–60), *Ciita* (59–61) and *Cxcl12* (62). Interestingly, we found IFI208, an interferon inducible gene, to be one of the most highly upregulated genes when HNRNPA2B1 was removed despite the overall dampening of IFN response signaling, suggesting a more complex role with a possible negative regulatory aspect for HNRNPA2B1. Likewise, we observed a downregulation of the costimulatory molecule CD86 both at the RNA and protein levels yet CD80, a costimulatory molecule with semi overlapping function, was increased suggesting specificity and complexity to the mechanisms by which HNRNPA2B1 regulates gene expression. It has been previously shown in an alternative septic shock model that macrophage CD80 plays a greater role in promoting inflammation than CD86 (63). Consistent with an alternative driver of protection, HNRNPA2B1 transcriptionally regulated expression of major components of the IFNG signaling pathway from the receptor *Ifngr* to the adaptor proteins, *JAK1,3* and *STAT1,3* as well as transcription factors (*IRF1,2,7,8,9*). Effects were not limited to the level of transcription and extended to reduced pathway activation through reduction in phosphorylation of STAT3 and the total STAT3 protein.

Given the changes we observed to IFN signaling and cytokine responses *ex vivo* upon loss of HNRNPA2B1, we chose to correlate those findings to *in vivo* challenge with LPS. Mice deficient in HNRNPA2B1 produced lower levels of proinflammatory cytokines including many produced predominantly by macrophages such as CCL2 (MCP1), CCL3 (MIP1a), CCL5 (Rantes), CXCL1 (KC) and IL6 in serum and spleen. This disruption in the inflammatory response proved advantageous to KO mice which did not become as hypothermic as the CTL mice following challenge. However, this attenuated inflammatory response proved detrimental to mice deficient in HNRNPA2B1 when infected with *Salmonella*. We chose *Salmonella* as it readily infects macrophages which provide a replicative niche that is abolished upon IFNG signaling and polarization of macrophages to an M1 state for pathogen clearance (42, 64, 65). Following *Salmonella* infection, the HNRNPA2B1 deficient mice succumbed earlier to infection and displayed higher bacterial burden, indicative of failure to clear the pathogen. This phenotype can be explained by the fact that the knockout mice have lower levels of the IFNG receptors, downstream adaptors, transcription factors, and co-stimulatory molecules. With all these pathways dampened, IFNG is no longer capable of coordinating cytokine activation (66–69) and sensitization of macrophages to potentiate a strong inflammatory response resulting in reduced effector cell (T cells and NK cells) activation and an inability to clear the pathogen (Fig. 6, H).

Mechanistically, HNRNPA2B1 -similar to other HNRNPs- is involved in RNA metabolism, specifically alternative splicing. We report its strict localization in the nucleus of macrophages at steady state and following stimulation with a variety of PRR ligands as well as Sendai and murine gamma herpesvirus (MHV) (Fig.5). A recent study by Wang et al. showed that HNRNPA2B1 acts as a DNA receptor and can translocate to the cytosol following HSV-1 infection, influencing IFN signaling through activation of TBK1-IRF3 (32). It is possible that movement of HNRNPA2B1 is stimulus dependent. Using RNA-seq we found that HNRNPA2B1 plays a role in the alternative splicing of the *Ifngr* locus. Macrophages deficient in HNRNPA2B1 showed an increase in an NGO isoform of *Ifngr* which lacks a start codon and is therefore not translated, and this was confirmed by lower expression of the IFNG receptor on the cell surface of HNRNPA2B1 knockout mice using flow cytometry. This downregulation of the IFNGR could explain the observed *in vivo* phenotypes as discussed earlier as lower expression of the receptor will dampen all the downstream responses following IFNG production impacting the overall adaptive immune response. Other IFN response genes such as *Stat1*, *Stat3*, *Irf7* and *Oas3* showed similar upregulation of an NGO transcript in the knockout macrophages, indicating HNRNPA2B1’s involvement in alternative splicing of other IFN response genes in the IFN pathway in a similar manner, thus regulating their expression pattern. Alternatively, some genes such as *Cd86* that are downregulated at the RNA and protein levels in the HNRNPA2B1 deficient mice could also be targets of alternative splicing, but they did not appear in our splicing analysis. It is technically difficult to capture transcripts such as NGO and other isoforms as they could be degraded more rapidly. Therefore, we do not know how many of the genes downregulated in the HNRNPA2B1 knockouts are controlled through splicing, transcription control or an alternative mechanism.

In conclusion, our findings highlight HNRNPA2B1’s integral role in promoting IFNG inflammatory responses in macrophages *in vivo*. Mechanistically HNRNPA2B1 ensures appropriate processing of the *IfngrI* transcript allowing adequate activation upon IFNG binding, driving expression of JAKs, STATs and IRF proteins as well as a large number of ISGs. This initiates macrophage activation and facilitates downstream activation of adaptive immunity. This work highlights HNRNPA2B1 as a key regulator of innate immunity providing new insights into how a ubiquitously expressed RNA binding protein can play unique roles in regulating IFN gene expression in macrophages.

## Materials and Methods

### Mice

Heterozygous floxed mice for the HNRNPA2B1 locus were generated using CRISPR to insert two loxP sites flanking exons 2 and 7 in the HNRNPA2B1 locus. Mice homozygous for the loxP sites were used as controls in these experiments. Heterozygous loxP mice were generated by Biocytogen where CRISPR/Cas9 was used to insert two loxP fragments in introns 1 and 7 of the HNRNPA2B1 locus. When exons 2-7 are removed a protein reading frame shift will occur which results in the production of a 100aa protein that eventually undergoes NMD.

#### CRISPR/Cas9 sgRNA

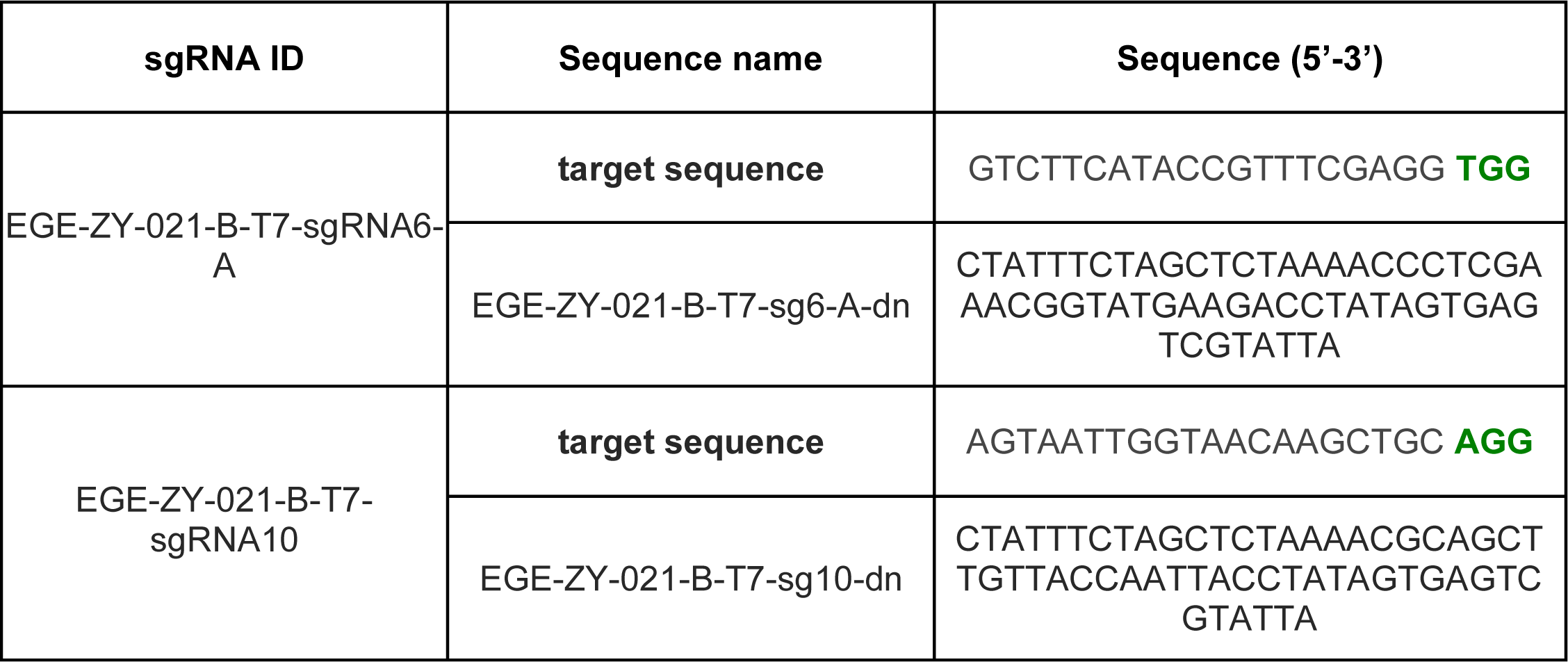

#### LoxP site integration detection primers

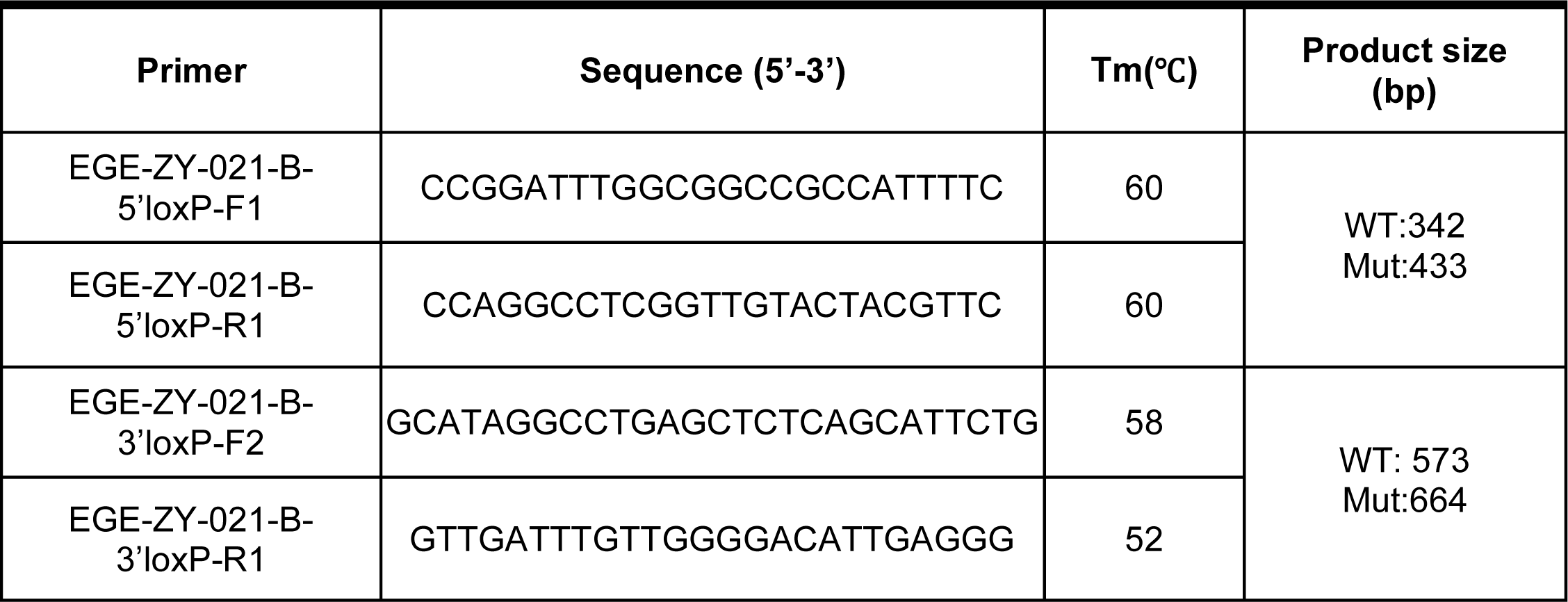

*LysMC*re mice were purchased from the Jackson Laboratory (Bar Harbor, ME), reference ID: RRID:IMSR_JAX:004781. Conditional Ko mice were homozygous for both the loxP sites and *LysM*Cre site. All mouse strains were bred at the University of California, Santa Cruz (UCSC) and maintained under specific pathogen-free conditions in the animal facilities of UCSC. All protocols were performed in accordance with the guidelines set forth by UCSC and Texas A&M Institutional Animal Care and Use Committees.

### Cell culture

BMDMs were generated by culturing erythrocyte-depleted BM cells in DMEM supplemented with 10% FCS, 5 mL pen/strep (100×), 500 μL ciprofloxacin (10 mg/mL), and 10% L929 supernatant for 7 to 14 d, with the replacement of culture medium every 2 to 3 d.

### *Ex vivo* stimulation of macrophages and inflammasome activation

Bone marrow derived macrophage cells were stimulated with Lipopolysaccharide (LPS) at 200 ng/ml (TLR4) for 5 hrs for RNA extraction and 18 hrs for ELISA and western blots. For inflammasome activation assay, cells were primed with either LPS at 20ng/mL or IFNγ at 50 ng/mL overnight followed by LPS for 3 hrs. Inflammasome was activated by ATP at 5mM for 2 hrs or dA:dT at 1ug/mL for 6 hrs or Nigericin at 25uM for 0.5hrs. For RNA and protein isolation, 1-2×10^6^cells were seeded in 12-well format or 10×10^6^ cells were seeded in 10cm plates.

Antibodies used for apoptosis assessment: Annexin V: Biolegend 640920 and PI.

### RNA isolation, cDNA synthesis and RT-qPCR

Total RNA was purified from cells or tissues using Direct-zol RNA MiniPrep Kit (Zymo Research, R2072) and TRIzol reagent (Ambion, T9424) according to the manufacturer’s instructions. RNA was quantified and assessed for purity using a nanodrop spectrometer (Thermo Fisher). Equal amounts of RNA (500 to 1,000 ng) were reverse transcribed using iScript Reverse Transcription Supermix (Bio-Rad, 1708841), followed by qPCR using iQ SYBR Green Supermix reagent (Bio-Rad, 1725122) with the following parameters: 95 °C for 10 min, followed by 40 cycles of 95 °C for 15 s, 60 °C for 30 s, and 72°C for 30s, followed by melt-curve analysis to control for nonspecific PCR amplifications. Oligos used in qPCR analysis were designed using Primer3 Input version 0.4.0.

Gene expression levels were normalized to *Actin* or *Hprt* as housekeeping genes.

### Cells Supernatant Collection for ELISA

Supernatant was collected from cultured and treated BMDMs, centrifuged at 12000xg, 5mins at RT and submitted for cytokine analysis.

### Serum Harvest

Mice were humanely sacrificed; blood was collected immediately postmortem by cardiac puncture. Blood was allowed to clot and centrifuged, serum was stored at −70°C, then sent to EVE for measurements of cytokines/chemokines.

### Spleen Tissue Harvesting for cytokine measurement

Mice were humanely sacrificed, and their spleens were excised. The whole spleens were snap frozen and homogenized, and the resulting homogenates were incubated on ice for 30 min and then centrifuged at 300 × *g* for 20 min. The supernatants were harvested, passed through a 0.45-μm-pore-size filter, and used immediately or stored at −70°C, then sent to EVE for measurements of cytokines/chemokines.

### RNA sequencing libraries

RNA-Seq was performed in BMDMs with no treatment or treated with LPS for 5 hrs. The data are accessible at the National Center for Biotechnology Information (NCBI) Gene Expression Omnibus (GEO) database, accession: GSE243269.

RNA-Seq was performed in biological triplicates in fl/fl BMDMs used as control and KO BMDMs at 0 and 5 h after LPS treatment (200 ng/mL). RNA-Seq libraries were generated from total RNA (1 μg) using the Bioo kit, quality was assessed, and samples were read on a High-SEq 4000 as paired-end 150-bp reads. Sequencing reads were aligned to the mouse genome (assembly GRCm38/mm10) using STAR. Differential gene-expression analyses were conducted using DESeq2. GO enrichment analysis was performed using PANTHER. Data was submitted to GEO, accession:GSE243269.

### Alternative Splicing Analysis

Alignment files from RNA-Seq analysis were filtered to include only canonical chromosomes and were passed to IUTA to test differential isoform usage. Gene isoforms from the Gencode M25 Comprehensive gene annotation file were used. Family wise error rate (FWER) was accounted for with Bonferroni correction and changes were called significant at FWER < 0.01.

### Cell Extracts and Western Blots

Cell lysates were prepared in RIPA buffer (150 mM NaCl, 1.0% Nonidet P-40, 0.5% sodium deoxycholate, 0.1% SDS, 50 mM Tris-HCl [pH 7.4], and 1.0 mM EDTA) containing protease-inhibitor mixture (Roche, 5892791001) and quantified using Pierce Bicinchoninic Acid Assay assay (Thermo Fisher, 23225). When indicated, the NEPER kit (Thermo Fisher Scientific, 78833) supplemented with protease inhibitor mixture (Roche) or 100 U/mL SUPERase-In (Ambion, AM2694) was used for cellular fractionation prior to Western blotting. Equivalent amounts (15 μg) of each sample were resolved by SDS-PAGE and transferred to polyvinylidene difluoride membranes using Trans-Blot Turbo Transfer System (Bio-Rad). Membranes were blocked with PBS, supplemented with 5% (wt/vol) nonfat dry milk for 1 h, and probed with primary antibodies overnight with either HNRNPA2B1 (1:1000, Santa Cruz Biotech, Sc-374053), pSTAT3 (1:500, Cell Signaling, 9138S), STAT3 (1:500, Cell Signaling, 9139S), pJAK3 (1:500, Cell Signaling, 5031S), or JAK3 (1:500, Cell Signaling, 8863S). Horseradish peroxidase-conjugated β-actin (1:500, Santa Cruz Biotechnology, sc-47778), HNRNPL (1:1000, Santa Cruz Biotech, Sc-32317) or GAPDH (1:1000, Santa Cruz Biotech, Sc-32233) were used as loading controls. Horseradish peroxidase-conjugated goat anti-mouse (1:10,000, Bio-Rad, #1721011) or anti-rabbit (1:2,000, Bio-Rad, #1706515) secondary antibodies were used. Western blots were developed using Amersham enhanced chemiluminescence (ECL) Prime chemiluminescent substrate (GE Healthcare, 45-002-401) or Pierce ECL (Life Technologies, 32106).

### *In Vivo* LPS-Induced Endotoxic Shock Assay

Age- and sex-matched CTL and KO mice (10 to 12 wk old) were i.p. injected with PBS as a control or *E. coli* LPS (5 mg/kg/animal). For gene expression and cytokine analysis, mice were euthanized 6 h or 18 h post injection. Blood was collected immediately postmortem by cardiac puncture. Serum was submitted to Eve Technologies for cytokine analysis.

### Assessment of Immune Cell Populations Using Flow Cytometry

Blood was collected immediately postmortem by cardiac puncture, and single-cell suspensions prepared from the spleen of CTL and HNRNPA2B1 KO mice were depleted of red blood cells (RBCs) prior to staining. Whole spleens were collected, homogenized and depleted of RBCs.

Fragment, crystallizable (Fc) receptors were blocked (anti-CD16/32, BD Pharmingen) prior to staining with LIVE/DEAD Fixable near IR Dead Cell Stain (Thermo Fisher), anti– CD11b-AlexaFluor 488, anti–LY6G-BV421, anti–LY6C-PE-cy7, anti–CD19 BV786, anti– CSF-1R APC, anti–CD3-PE, anti–SiglecF Super Bright 645, and anti–F4/80 PE-eFluor 610, MHCII BV421, CD86 Super Bright 645, CD80 PE-Cy7, IFNGRI BV605, IFNGRII PE, all by Thermo Fisher.

Flow cytometry analysis of BMDMs harvested from CTL and HNRNPA2B1 KO mice was performed; cells were stained with LIVE/DEAD Fixable near IR Dead Cell Stain (Thermo Fisher),anti–CD11b-AlexaFluor 488, anti–F4/80 PE-eFluor 610, MHCII BV421, CD86 Super Bright 645, CD80 PE-Cy7, IFNGRI BV605, IFNGRII PE, all by Thermo Fisher.

Data acquisition was performed using Attune NxT (Thermo Fisher). Analysis was performed using FlowJo analysis software (BD Biosciences).

### Proliferation Assay

CTL and KO BMDMs were cultured as described above. Cell proliferation kit (MTT) by Millipore Sigma was used to assess cell proliferation. Solubilization of the purple formazan crystals was measured as an indicator of metabolically active cells through spectrophotometric absorbance of the samples using a microplate (ELISA) reader.

### Phagocytosis Assay

CTL and KO BMDMs were cultured as described above. Cells were then administered with pHrodo Green *E. coli* BioParticles (Invitrogen) conjugates for 0, 15, 30, and 60 min at 37 °C. pHrodo green fluorescence, a measure of *E. coli* in an acidified phagosome, was examined by flow cytometry.

### *S. enterica* (ser. Typhimirium)

*Salmonella enterica* serovar Typhimurium (SL1344) was obtained from Dr. Denise Monack, Stanford. S. T. stocks were streaked out on LB agar plates and incubated at 37 °C overnight. For S. Typhimurium infection, overnight cultures of bacteria were grown in LB broth containing 0.3 M NaCl and grown at 37 °C until they reached an OD600 of 0.9. On the day of infection cultures were diluted 1:20. Once cultures had reached mid-log phase (OD600 0.6-0.8) 2-3 hrs, 1 mL of bacteria were pelleted at 5000 rpm for 3 min and washed twice with 2X PBS.

### *In vivo* S. enterica (ser. Typhimurium) infection

Age- and sex-matched CTL and KO mice (10 to 12 wk old) were i.p. injected with *Salmonella enterica* serovar Typhimurium 2.5 x 10^4^ bugs per animal. For CFUs and cytokine analysis, mice were euthanized 3 days post infection. Blood was collected immediately postmortem by cardiac puncture. Serum was submitted to Eve Technologies for cytokine analysis. For time to death mice were monitored for signs of imminent morbidity (e.g. hunched posture, ruffled coat, lethargy as described in <u>22822473</u>) while blinded to genotype.

## KEY RESOURCES TABLE

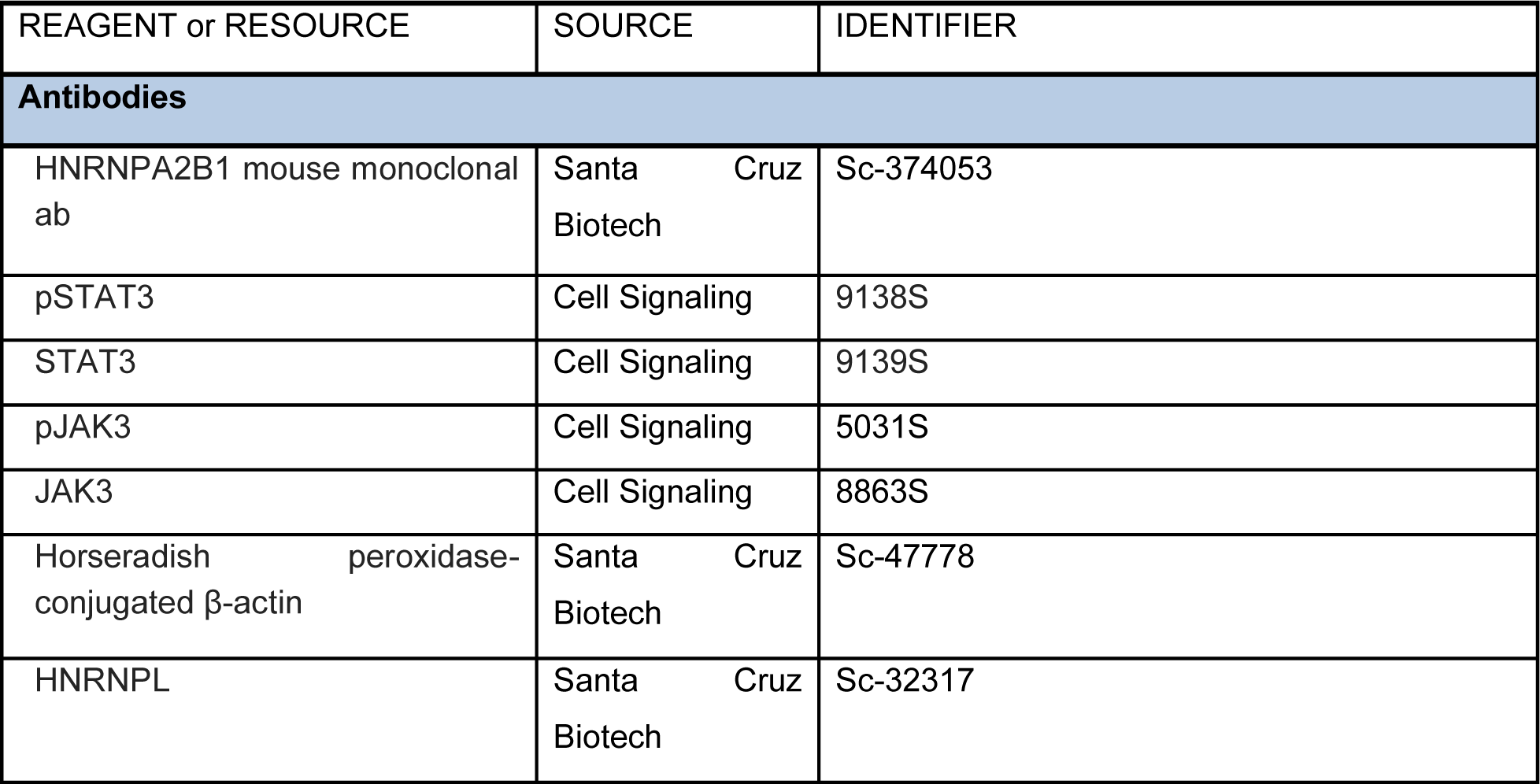
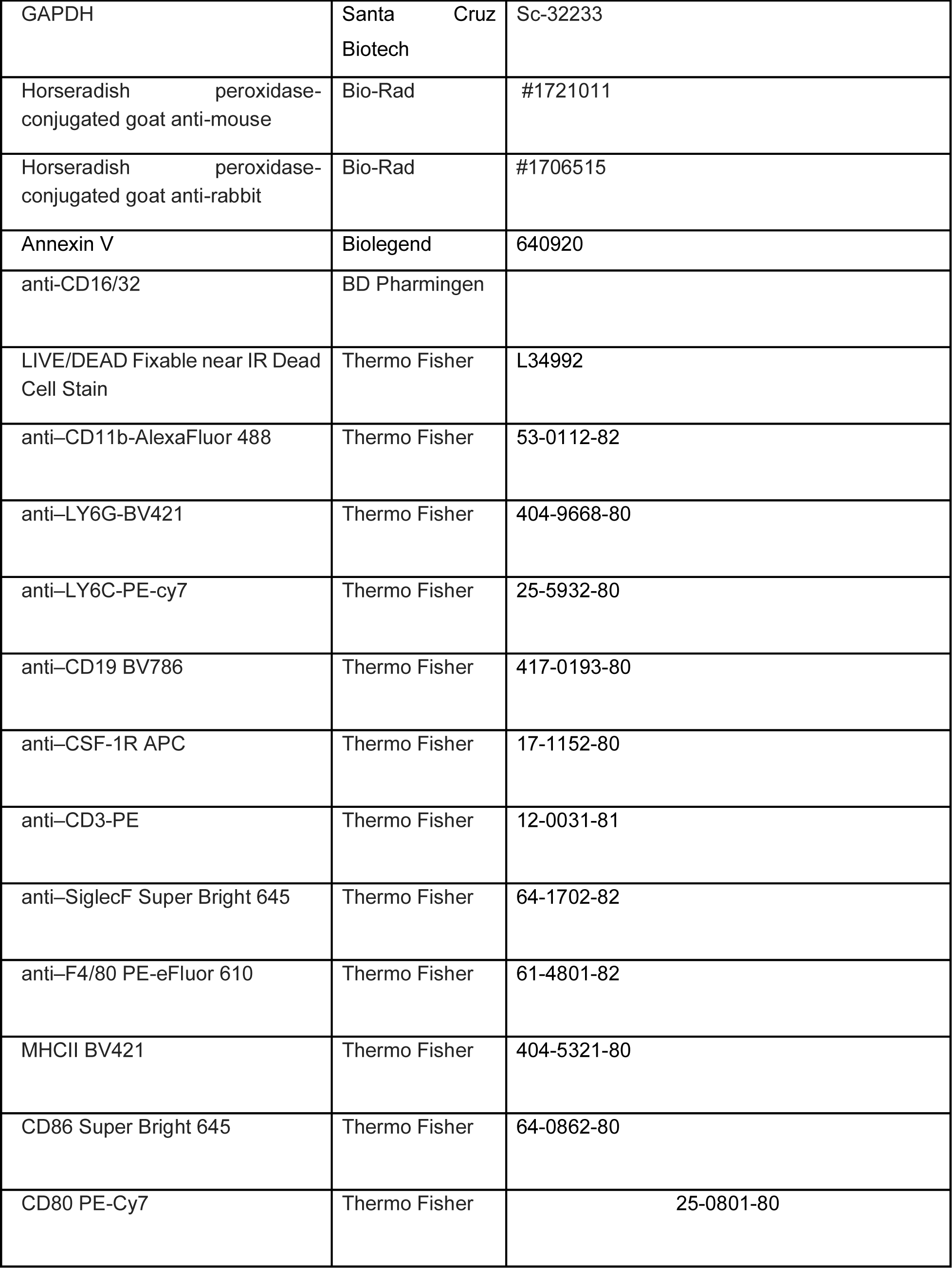
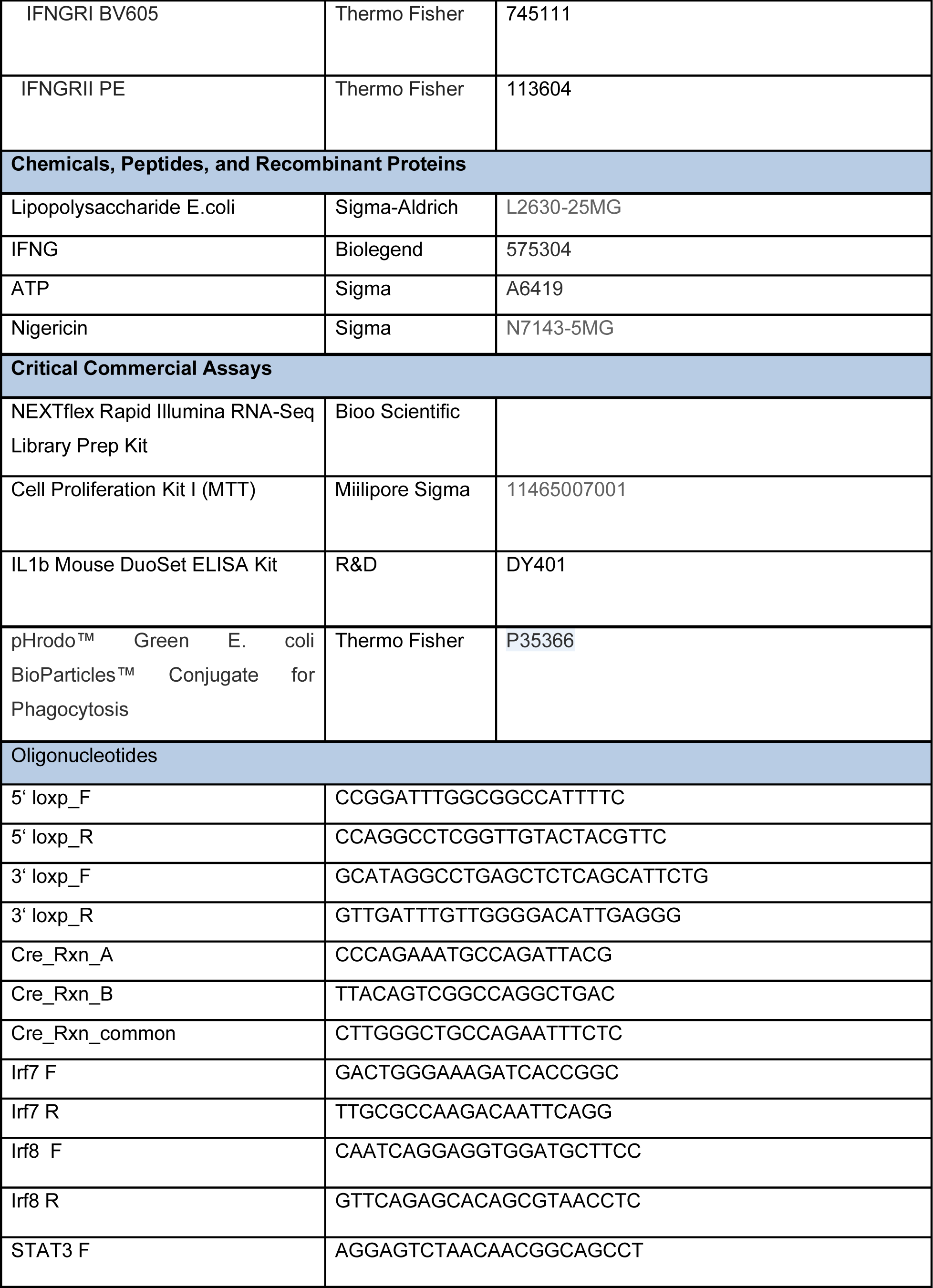
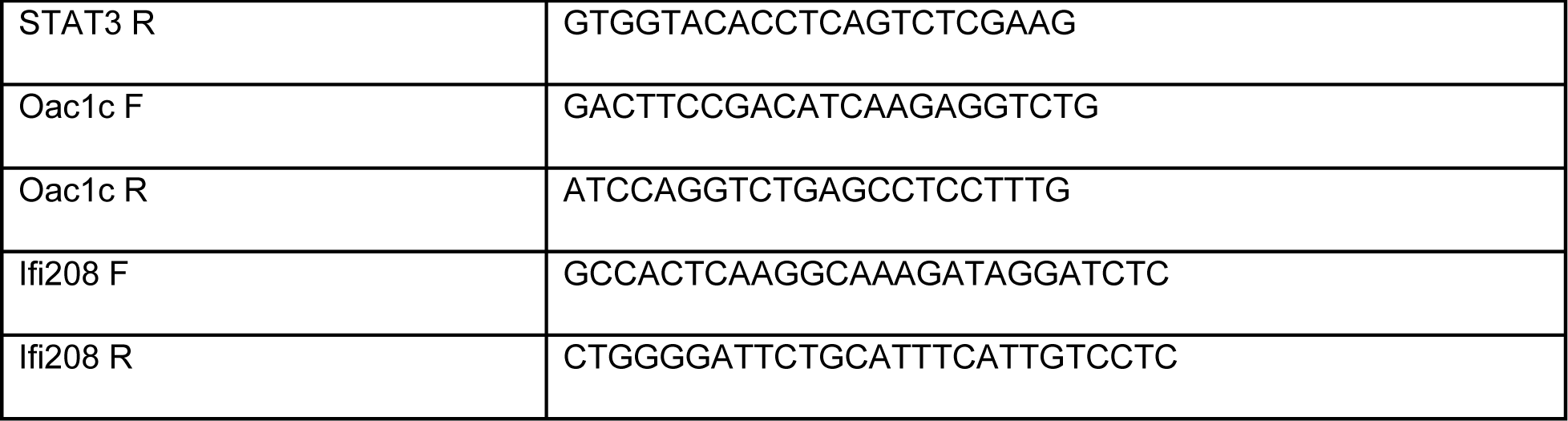

## Acknowledgements

SC is supported by R01 AI148413 from NIAID and R35GM137801 from NIGMS. MMS is supported by California Institute of Regenerative Medicine/ IBSC predoctoral training program. We would like to thank Dr. Angela Brooks for her valuable feedback on the splicing aspects of this manuscript. Graphical abstract was generated using Biorender.com.

## Author contributions

SC and MMS conceptualized and acquired funding for this study. MMS performed all *ex vivo* experiments and *in vivo* endotoxic shock studies as well as data analysis. CW, CM, AC and SA performed *Salmonella* infections and harvested all samples for study. SK performed RNA-sequencing analysis and EM performed alternative splicing analysis for this study. MMS wrote the initial draft of this manuscript. All authors contributed to reviewing and editing of the manuscript.

## Competing interests

S.C is a paid consultant to NextRNA Therapeutics.

---

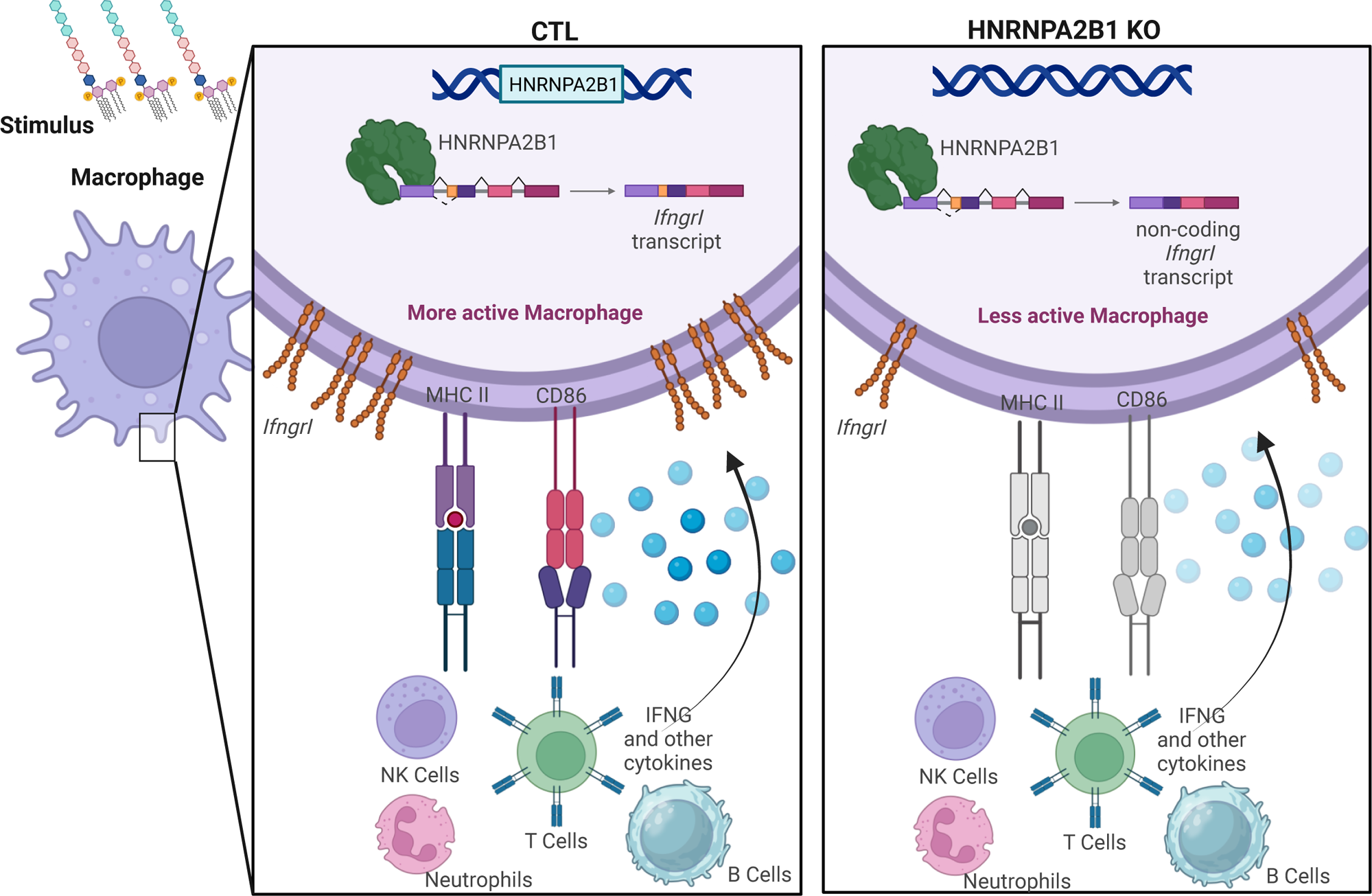

