## Supplemental files for "The RNA binding protein, HNRNPA2B1, regulates IFNG signaling in macrophages"

### Figure legends

**Supplementary figure 1. Genotyping HNRNPA2B1 KO mouse. A.** Primer design for primer sets A and B which identify successful integration of the loxP sites flanking exons 2 and 7. **B.** Genotyping gel showing band sizes for HNRNPA2B1 locus as CTL, heterozygous for loxP and homozygous for loxP. **C.** Primer design to identify successful integration of cre recombinase downstream *Lyz2* promoter. **D.** Genotyping gel showing band sizes for CTL, Cre heterozygous and Cre homozygous locus.

**Supplementary figure 2. HNRNPA2B1 regulates IFN response genes transcriptionally. A-E.** Normalized qRTPCR results for **A.** *Irf7***, B.** *Irf8***, C.** *Stat3***, D.** *Oas1c* and **E.** *Ifi208* Showing changes in CTL and KO BMDMs at baseline and after LPS stimulation. Student's t-tests were performed using GraphPad Prism. Asterisks indicate statistically significant differences between mouse lines (*P ≤ 0.05).

**Supplementary figure 3. HNRNPA2B1 regulates inflammatory cytokine production under stimulus. A-M.** Multiplex ELISA results of inflammatory cytokine changes in CTL and KO BMDM supernatant after treatment with LPS for 18 hrs. **A**. MCSF, **B.** GCSF, **C.** TARC, **D.** IL1B, **E.** IL5, **F.** IL12, **G.** Il15, **H.** TIMP1, **I.** MCP-1, **J.** MCP-5, **K.** Il16, **L.** IFNG, **M.** VEGF. Student's t-tests were performed using GraphPad Prism. Asterisks indicate statistically significant differences between mouse lines (*P ≤ 0.05, **P ≤ 0.01).

**Supplementary figure 4. Immune cell profiling in blood and spleen of CTL and KO mice. A-D.** Immune cell levels in CTL and KO mice blood analyzed using flow cytometry **A.** T cells**, B.** B cells**, C.** Monocytes**, D.** Eosinophils**. E-H.** Immune cell levels in CTL and KO mice spleen analyzed using flow cytometry **E.** T cells**, F.** B cells**, G.** Monocytes**, H.** Eosinophils**. I.** Levels of small peritoneal macrophages**, J.** Levels of large peritoneal macrophages.

**Supplementary figure 5. HNRNPA2B1 remains in the nucleus at baseline and after exposure to stimulus. A.** Immunofluorescence Images of primary BMDMs. Nuclei stained with DAPI, HNRNPA2B1 stained with an anti-mouse secondary antibody conjugated to Alexa 488. Image processing using Fiji, nuclei boundaries were drawn using masks on DAPI channel and overlaid on to green channel to show that HNRNPA2B1 exists exclusively in the nucleus.

**Supplementary figure 6. HNRNPA2B1 regulates alternative splicing of interferon response genes. A-D.** Isoform usage levels analyzed by IUTA showing switching in isoform usage between CTL and KO BMDMs in **A.** *Stat3***, B.** *Stat1***, C.** *Irf7* **and D.** *Oas3***.**

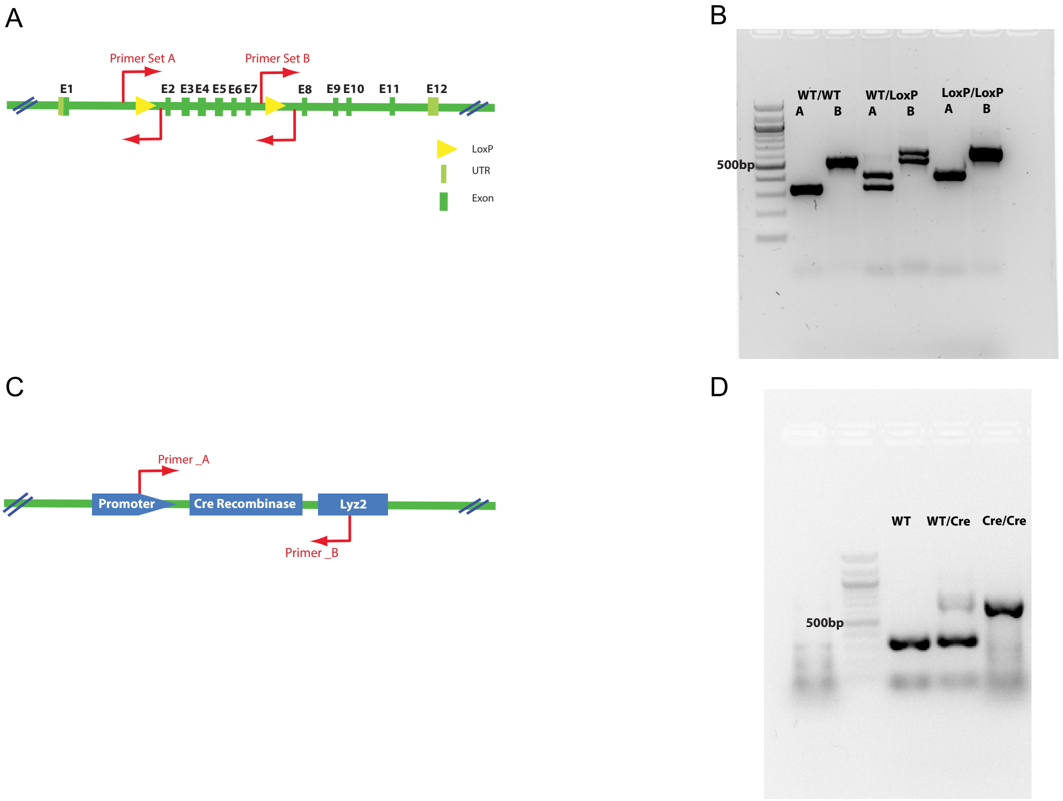
A
Primer Set A
Primer Set B
E1
E2 E3 E4 E5 E6 E7
E8
E9 E10
E11
E12
LoxP
UTR
Exon
B
WT/WT
A B
WT/LoxP
A B
LoxP/LoxP
A B
500bp
C
Primer _A
Promoter
Cre Recombinase
Lyz2
Primer _B
D
WT
WT/Cre
Cre/Cre
500bp

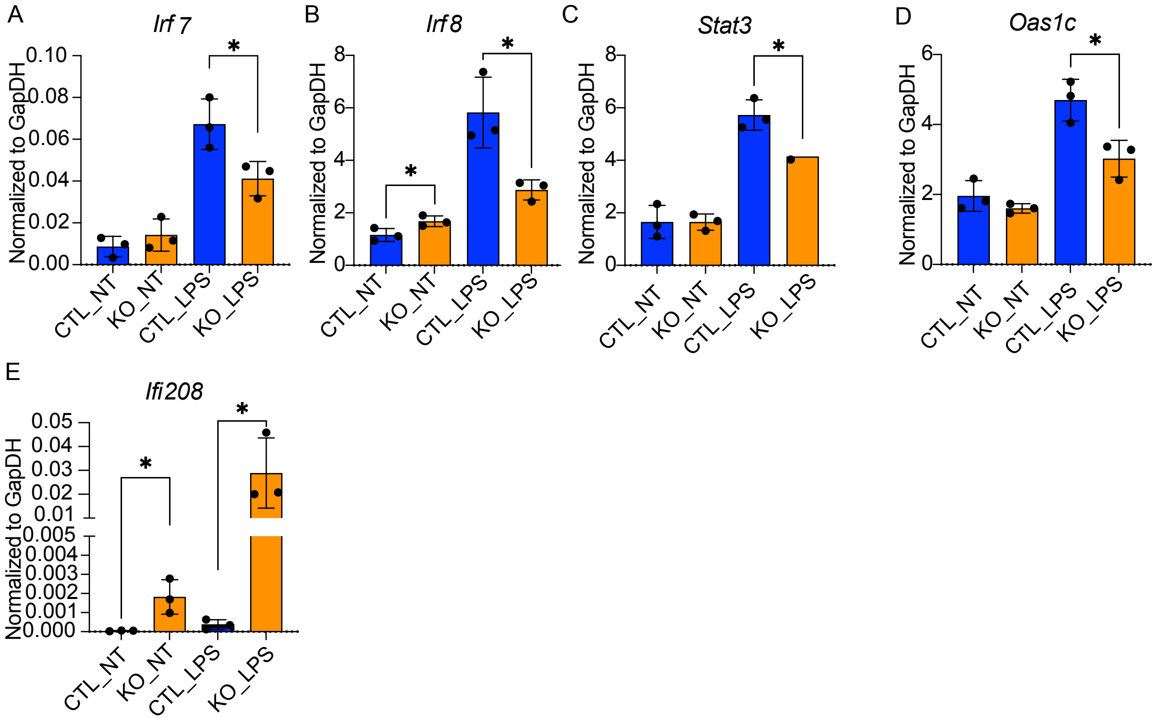

A
Irf 7
Normalized to GapDH
0.10
0.08
0.06
0.04
0.02
0.00
*
CTL_NT
KO_NT
CTL_LPS
KO_LPS
B
Irf 8
Normalized to GapDH
8
6
4
2
0
*
*
CTL_NT
KO_NT
CTL_LPS
KO_LPS
C
Stat3
Normalized to GapDH
8
6
4
2
0
*
CTL_NT
KO_NT
CTL_LPS
KO_LPS
D
Oas1c
Normalized to GapDH
6
4
2
0
*
CTL_NT
KO_NT
CTL_LPS
KO_LPS
E
Ifi208
Normalized to GapDH
0.05
0.04
0.03
0.02
0.01
0.005
0.004
0.003
0.002
0.001
0.000
*
*
CTL_NT
KO_NT
CTL_LPS
KO_LPS

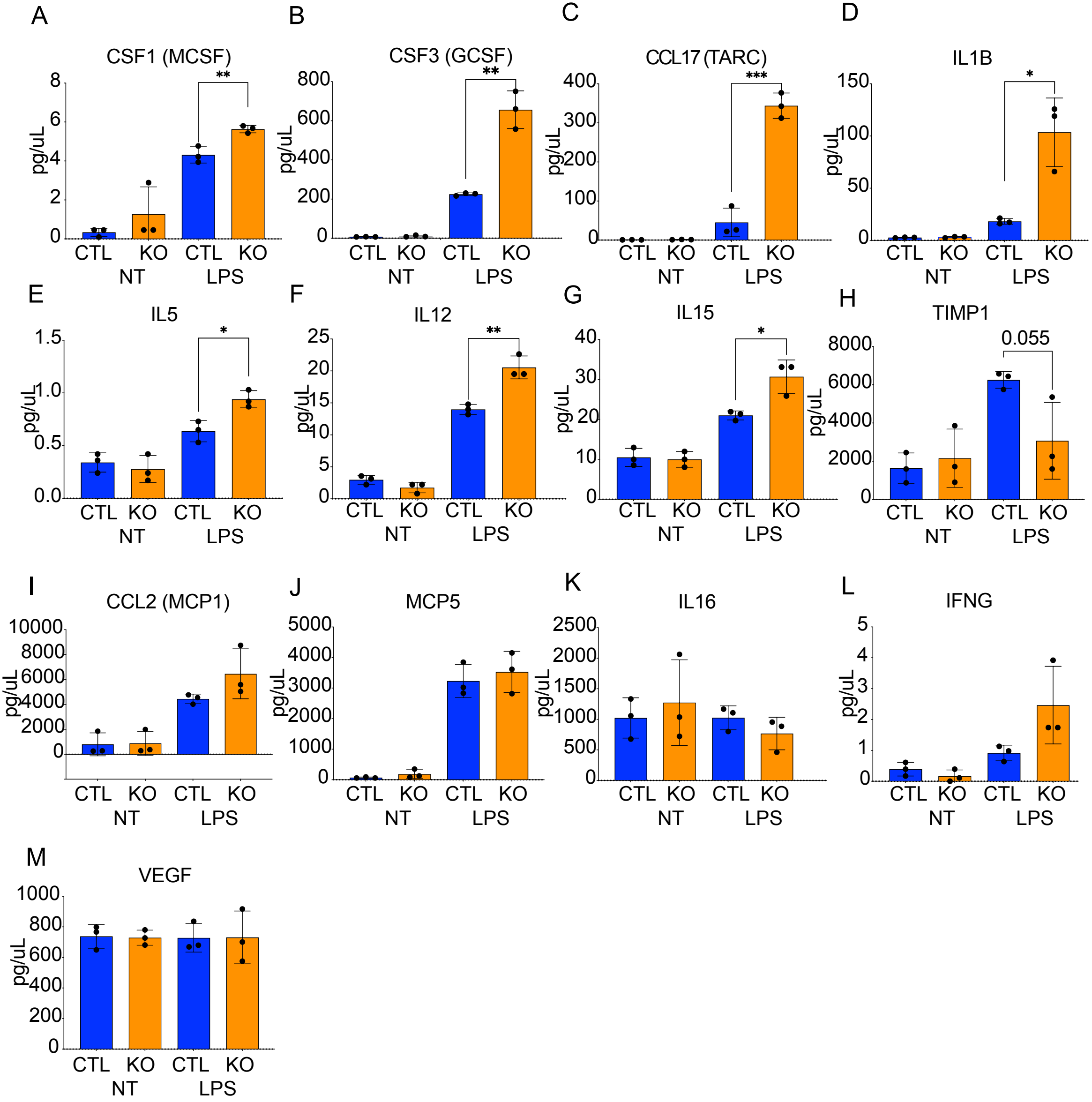

A
CSF1 (MCSF)
**
pg/uL
8
6
4
2
0
CTL KO CTL KO
NT LPS
B
CSF3 (GCSF)
**
pg/uL
800
600
400
200
0
CTL KO CTL KO
NT LPS
C
CCL17 (TARC)
***
pg/uL
400
300
200
100
0
CTL KO CTL KO
NT LPS
D
IL1B
*
pg/uL
150
100
50
0
CTL KO CTL KO
NT LPS
E
IL5
*
pg/uL
1.5
1.0
0.5
0.0
CTL KO CTL KO
NT LPS
F
IL12
**
pg/uL
25
20
15
10
5
0
CTL KO CTL KO
NT LPS
G
IL15
*
pg/uL
40
30
20
10
0
CTL KO CTL KO
NT LPS
H
TIMP1
0.055
pg/uL
8000
6000
4000
2000
0
CTL KO CTL KO
NT LPS
I
CCL2 (MCP1)
pg/uL
10000
8000
6000
4000
2000
0
CTL KO CTL KO
NT LPS
J
MCP5
pg/uL
5000
4000
3000
2000
1000
0
CTL KO CTL KO
NT LPS
K
IL16
pg/uL
2500
2000
1500
1000
500
0
CTL KO CTL KO
NT LPS
L
IFNG
pg/uL
5
4
3
2
1
0
CTL KO CTL KO
NT LPS
M
VEGF
pg/uL
1000
800
600
400
200
0
CTL KO CTL KO
NT LPS

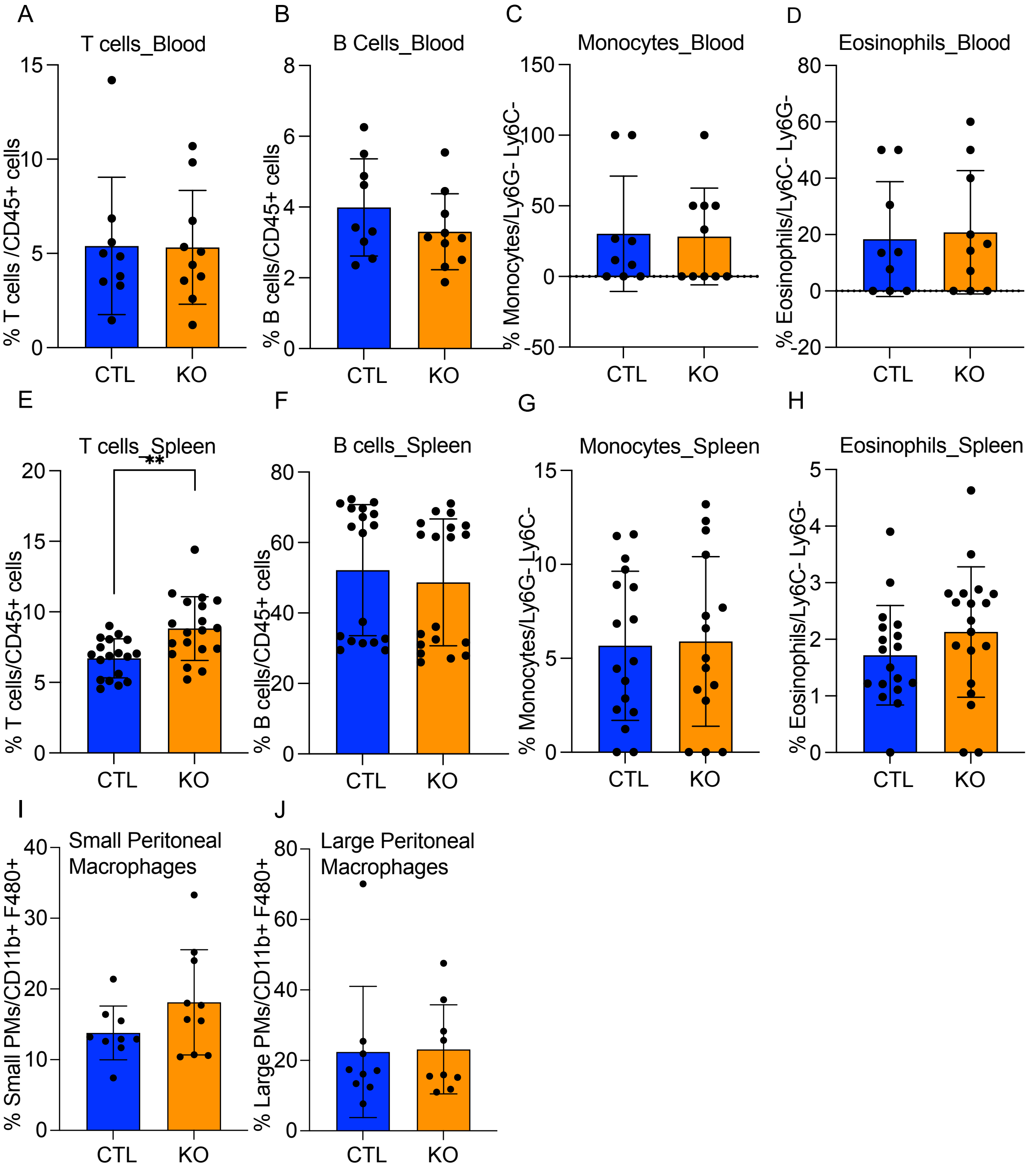
A
T cells_Blood
% T cells /CD45+ cells
B
B Cells_Blood
% B cells/CD45+ cells
C
Monocytes_Blood
% Monocytes/Ly6G- Ly6C-
D
Eosinophils_Blood
% Eosinophils/Ly6C- Ly6G-
E
T cells_Spleen
**
% T cells/CD45+ cells
F
B cells_Spleen
% B cells/CD45+ cells
G
Monocytes_Spleen
% Monocytes/Ly6G- Ly6C-
H
Eosinophils_Spleen
% Eosinophils/Ly6C- Ly6G-
I
Small Peritoneal Macrophages
% Small PMs/CD11b+ F480+
J
Large Peritoneal Macrophages
% Large PMs/CD11b+ F480+
CTL
KO

A

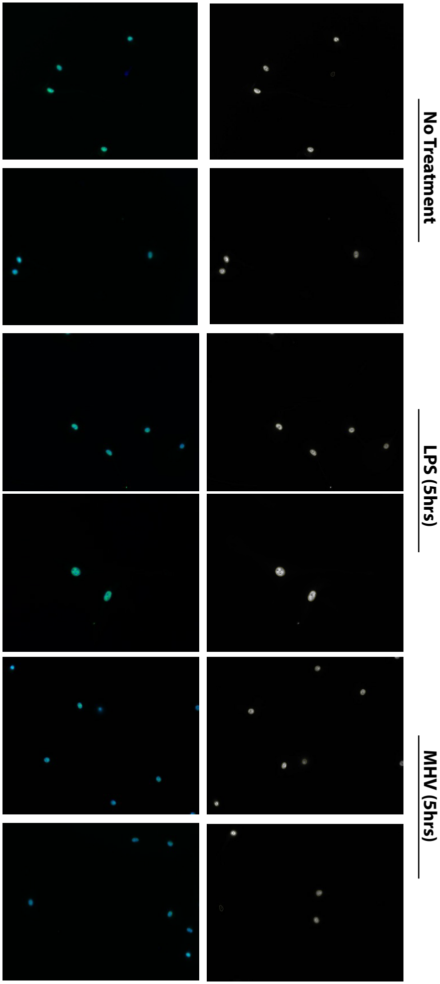

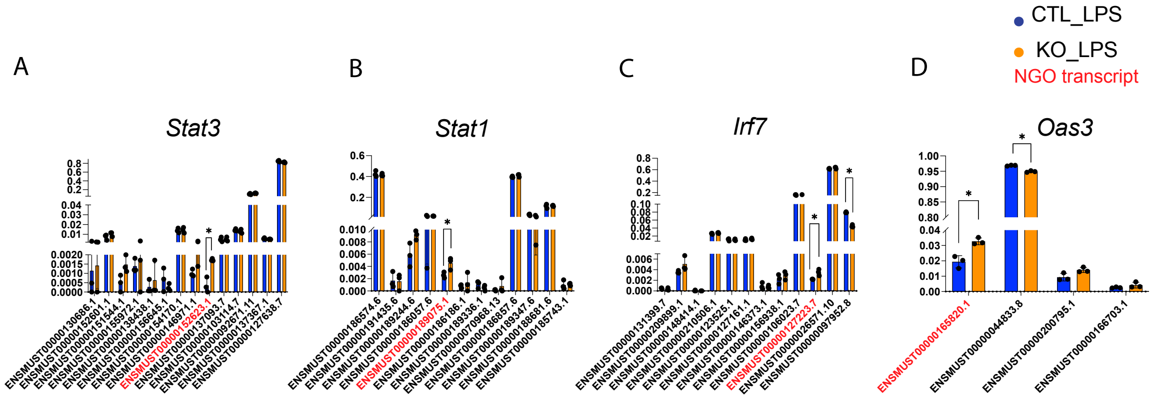
CTL_LPS
KO_LPS
NGO transcript
A
Stat3
ENSMUST00000130686.1
ENSMUST00000152601.1
ENSMUST00000151544.1
ENSMUST00000155972.1
ENSMUST00000138438.1
ENSMUST00000156645.1
ENSMUST00000154170.1
ENSMUST00000146971.1
ENSMUST00000152623.1
ENSMUST00000137093.7
ENSMUST00000103114.7
ENSMUST00000092671.11
ENSMUST00000137367.1
ENSMUST00000127638.7
B
Stat1
ENSMUST00000186574.6
ENSMUST00000191435.6
ENSMUST00000189244.6
ENSMUST00000186057.6
ENSMUST00000189075.1
ENSMUST00000186186.1
ENSMUST00000189336.1
ENSMUST00000070968.13
ENSMUST00000186857.6
ENSMUST00000189347.6
ENSMUST00000188681.6
ENSMUST00000185743.1
C
Irf7
ENSMUST00000131399.7
ENSMUST00000209899.1
ENSMUST00000148414.1
ENSMUST00000210506.1
ENSMUST00000123525.1
ENSMUST00000127161.1
ENSMUST00000146373.1
ENSMUST00000156938.1
ENSMUST00000106023.7
ENSMUST00000127223.7
ENSMUST00000026571.10
ENSMUST00000097952.8
D
Oas3
ENSMUST00000165820.1
ENSMUST00000044833.8
ENSMUST00000200795.1
ENSMUST00000166703.1
